# Cell competition corrects noisy Wnt morphogen gradients to achieve robust patterning

**DOI:** 10.1101/423814

**Authors:** Yuki Akieda, Shohei Ogamino, Hironobu Furuie, Shizuka Ishitani, Ryutaro Akiyoshi, Jumpei Nogami, Takamasa Masuda, Nobuyuki Shimizu, Yasuyuki Ohkawa, Tohru Ishitani

**Affiliations:** Laboratory of Integrated Signaling Systems, Department of Molecular Medicine, Institute for Molecular & Cellular Regulation, Gunma University, Gunma, 371-8512, Japan; Medical Institute of Bioregulation, Kyushu University, Fukuoka 812-8582, Japan; Graduate School of Medical Sciences, Kyushu University, Fukuoka 812-8582, Japan; Evaluation Technology Department 1, Medical R&D Planning Division, Olympus Corp., Tokyo, 192-8512, Japan; Division of Transcriptomics, Medical Institute of Bioregulation, Kyushu University, Fukuoka 812-8582, Japan

**Keywords:** Morphogen gradient, Wnt/β-catenin signaling, cell competition, tissue patterning, robustness

## Abstract

Morphogen signaling forms an activity gradient and instructs cell identities in a signaling strength-dependent manner to pattern developing tissues. However, developing tissues also undergo dynamic morphogenesis, which may produce cells with unfit morphogen signaling and consequent noisy morphogen gradient. Here we show that a cell competition-related system corrects such noisy morphogen gradients. Zebrafish imaging analyses of the Wnt/β-catenin signaling-gradient, which acts as a morphogen to establish embryonic anterior-posterior patterning, revealed that unfit cells with abnormal Wnt/β-catenin activity spontaneously appear and produce noise in the Wnt/β-catenin-gradient. Communication between the unfit and neighboring fit cells via cadherin proteins stimulates the apoptosis of the unfit cells by activating Smad signaling and reactive oxygen species production. This unfit cell elimination is required for proper Wnt/β-catenin-gradient formation and consequent anterior-posterior patterning. Because this gradient controls patterning not only in the embryo but also in adult tissues, this system may support tissue robustness and disease prevention.

## INTRODUCTION

Tissue patterning is a fundamental process during embryonic development and adult tissue homeostasis. To reproducibly achieve precise tissue patterning, the molecular and cellular systems controlling patterning must be robust against environmental and physiological perturbations. Activity gradients of morphogen signaling, including Wnt/β-catenin, bone morphogenetic protein (BMP), sonic hedgehog (Shh), fibroblast growth factor (FGF), and nodal signaling pattern the tissue axes (Schier and Talbot, 2005; Wolpert, 1969). In the embryonic anterior-posterior (AP) axis formation of deuterostomes (amphioxus, fish, frog, and mammal), Wnt/β-catenin signaling is activated and inhibited in the presumptive posterior and anterior tissue, respectively. This bi-directional regulation forms a signaling activity gradient along the AP axis to establish embryonic AP patterning (Petersen and Reddien, 2009). However, rapid cell proliferation and movement in developing tissues may affect morphogen diffusion and signal transduction and produce cells with unfit signaling and consequent noisy morphogen gradients. How these noises are overcome to generate robust patterning remains incompletely understood.

Cell competition is an interactive process wherein cells compete for fitness in a tissue environment, with relatively higher fitness cells eliminating those with lower fitness (Merino et al., 2016; Morata and Ripoll, 1975). Key features include originating from specific interactions between two cell types leading to elimination in one, and context-dependency. Mosaically introduced polarity-deficient cells are apoptotically eliminated by their neighboring wild-type cells in *Drosophila* imaginal disc and mammalian cultured cells (Brumby and Richardson, 2003; Grzeschik et al., 2007; Norman et al., 2012) and in Myc-low-level cells upon communicating with Myc-high-level cells (Clavería et al., 2013; Johnston et al., 1999; Sancho et al., 2013). Although cell competition, evolutionarily conserved from insects to mammals, may assist in proper embryogenesis, tissue morphogenesis, and tumor progression and prevention (Di-Gregorio et al., 2016), its physiological relevance and detailed mechanisms, especially of unfit cell-sensing, remain unclear.

Here, we reveal a cell competition-related system for correcting the noise in the Wnt/β-catenin morphogen gradient, presenting a new physiological role of cell competition and the mechanisms that mediate unfit cell sensing and elimination.

## RESULTS

### Apoptotic Elimination of Unfit Cells Smoothens the Wnt/β-catenin Gradient

To clarify the entire morphogen gradient formation process, we visualized Wnt/β-catenin signaling activity during AP axis formation in zebrafish early embryos (Figure 1A) using OTM:d2EGFP (Shimizu et al., 2012) and OTM:Eluc-CP (Figure S1A) reporters. These respectively express destabilized EGFP (d2EGFP), providing high spatial resolution, and highly-destabilized Emerald luciferase (Eluc-CP), possessing high temporal resolution and suitable for quantitative analyzes (Figures S1B–E), upon Wnt/β-catenin signaling activation. A noisy signaling-gradient along the AP axis was detected in both transgenic zebrafish embryo types at around 8.5–12 hours-post-fertilization (hpf) (Figures 1B–D, Movie S1). Abnormally low- and high- Wnt/β-catenin activities were spontaneously detected in the Wnt/β-catenin activity-high posterior and -low regions, respectively (Figures 1B, 1D, 1E, Movie S1). Unfit cells with abnormal Wnt/β-catenin activity gradually disappeared over time (Figure 1D), suggesting that zebrafish embryonic tissue may possess a system for eliminating signaling noise to smoothen the Wnt/β-catenin-gradient. As mouse embryonic tissues eliminate defective cells, including low Myc level, autophagy-deficient, and tetraploid cells in an apoptosis-dependent manner (Clavería et al., 2013; Sancho et al., 2013), unfit Wnt/β-catenin activity-abnormal cells might also be apoptotically eliminated. To investigate the relationship between apoptosis and abnormal Wnt/β-catenin activity, we detected active caspase-3- and TUNEL-positive apoptotic cells in 8–10 hpf embryos undergoing Wnt/β-catenin-gradient-mediated AP axis patterning (Figures S2A–C, Movie S2). Apoptotic cell number and position varied between embryos (Figures S2D and S2E), suggesting that the apoptosis is not pre-programmed. In some unfit Wnt/β-catenin activity-abnormal cells, caspase-3 was activated (Figure 1B, right), whereas apoptosis inhibition by anti-apoptotic bcl-2 or caspase inhibitor p35 overexpression reduced physiologically occurring apoptosis (Figures S2C and S2F), enhanced the appearance of unfit cells with abnormally high or low Wnt/β-catenin activity, and severely distorted the Wnt/β-catenin activity gradient (Figures 1F and S2G). These results suggest that apoptotic elimination of unfit cells smoothens the Wnt/β-catenin-gradient.

**Figure 1.**
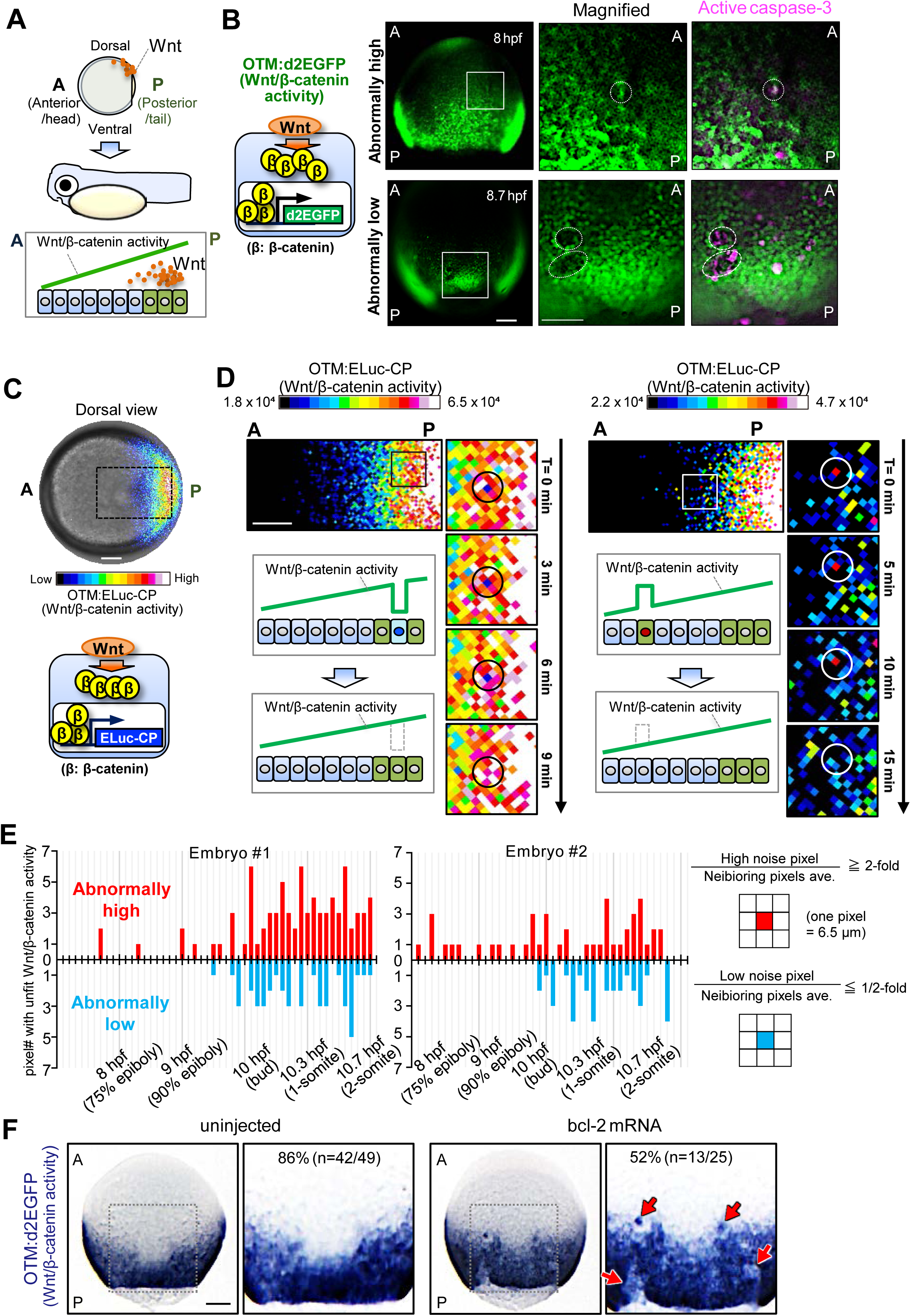
Apoptotic Elimination of Unfit Cells Smoothens the Wnt/β-Catenin Gradient. (A) Schematic illustration of Wnt/β-catenin activity gradient formation. A: anterior, P: posterior. (B) Caspase-3 activation in unfit cells with abnormal Wnt/β-catenin activity. Whole-mount immunostaining of d2EGFP (green) and active caspase-3 (magenta) in Tg(OTM:d2EGFP) zebrafish embryos (Dorsal view). Dotted line indicates abnormal Wnt/β-catenin-reporter activity. Scale bars, 200 μm. (C) OTM:ELuc-CP drives destabilized ELuc-CP expression in response to Wnt/β-catenin signaling activation in reporter embryo (dorsal view). Scale bar, 200 μm. (See also Movie S1). (D) Time lapse images showing unfit cells with abnormal Wnt/β-catenin activity appear then disappear. Scale bars, 100 μm. Pixel area length is 6.5 μm, ≤ zebrafish deep cell diameter (~10 μm). (E) Physiological Wnt/β-catenin-noise during zebrafish AP axis formation. Graphs show the number of pixels with unfit Wnt/β-catenin activity in the luminescence images of living OTM:ELuc-CP transgenic zebrafish embryos during AP axis formation. Schematic illustrations: pixel retaining > 2-fold or < 2-fold intensity compared to neighboring pixels for ≥ frames (> 6 min) was defined as High or Low noise, respectively. Pixels retained for ≥ 2 frames were counted as the physiological Wnt/β-catenin noise. Pixels spontaneously showing abnormally high or low activity within one frame were regarded as “other noise” (e.g., cosmic rays and detector noises) and excluded. (F) Apoptosis inhibition enhances unfit abnormally-high or-low Wnt/β-catenin activity cell appearance. Whole-mount *in situ* hybridization of d2EGFP in Tg(OTM:d2EGFP) embryos (dorsal view). Scale bar, 200 μm. (Statistical details and sample numbers are described in Table S1). See also Figures S1 and S2.

### Substantial Difference in Wnt/β-catenin Activity Between Unfit and Neighboring Cells Triggers Unfit Cell Apoptosis

To confirm that embryonic tissue equips the system for eliminating cells with unfit Wnt/β-catenin activity via apoptosis, we artificially introduced a small number of GFP-expressing Wnt/β-catenin-abnormal cells into zebrafish embryos by injecting heat-shock-driven expression plasmids (Figure 2A) and tracking their behavior. As predicted, Wnt/β-catenin-hyperactivated cells expressing constitutively active β-catenin (β-catCA), a Wnt receptor LRP6 (LRP6CA), or a dominant-negative Wnt/β-catenin negative regulator GSK-3β mutant (GSK-3β DN) activated caspase-3 and gradually disappeared, whereas negative control cells expressing GFP alone survived. Wnt/β-catenin-inactivated cells expressing Wnt/β-catenin negative regulators Axin1 or GSK-3β or a dominant-negative LRP6 mutant also activated caspase-3 and were eliminated (Figures 2B-D, Movies S3). The Wnt/β-catenin-hyperactivated or -inactivated cells underwent DNA fragmentation (Figure S3A) and were TUNEL-positive (Figure S3B). Caspase-inhibitor p35 co-expression blocked β-catCA-expressing cell elimination from larval tissue (Figure 2E). These results suggest that the artificially introduced Wnt/β-catenin-abnormal cells are apoptotically eliminated.

**Figure 2.**
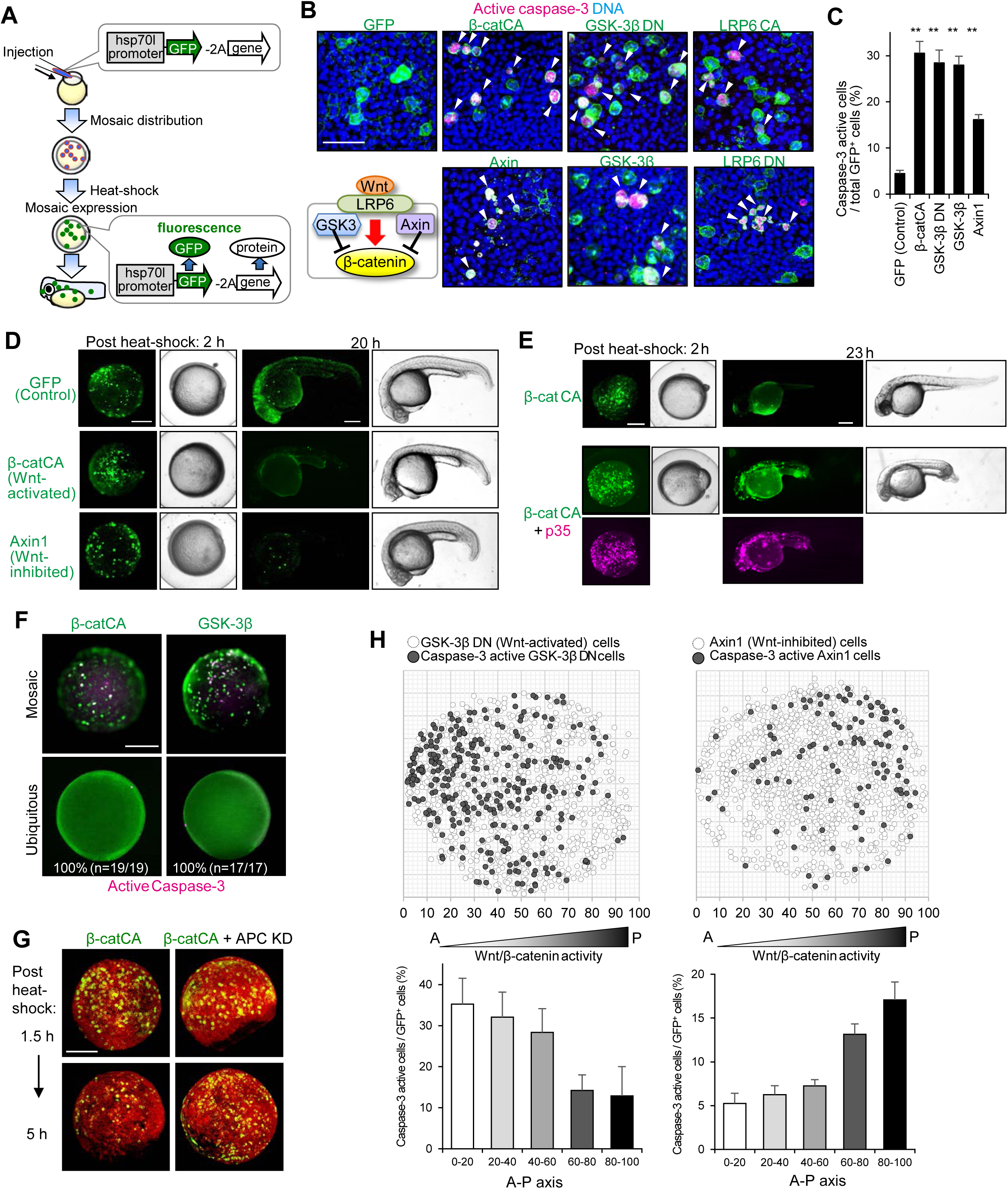
Substantial Difference in Wnt/β-catenin Activity Between Unfit and Neighboring Cells Triggers Unfit Cell Apoptosis. (A) Schematic illustration of experimental introduction of fluorescent Wnt/β-catenin-abnormal cells in zebrafish early embryo through heat shock induction. (B, C) Artificially introduced Wnt/β-catenin-abnormal cells undergo apoptosis. Confocal microscopy images showing whole-mount immunostaining of mosaic embryos expressing membrane GFP +/− Wnt activators or inhibitors. Scale bar, 50 μm. (C) Means ± SEM of GFP^+^ caspase-3-active cell frequencies. \*\**p* < 0.01. (D) Artificially introduced Wnt/β-catenin-abnormal cell disappearance from embryonic tissues. Scale bar, 200 μm. (See also Movies S3 and S4). (E) Embryos artificially introduced with cells expressing membrane GFP alone (GFP) or with β-catCA with or without caspase inhibitor p35. Fluorescence and bright-field images after heat-shock. p35 expression (magenta). Scale bar, 200 μm. (F) Surrounding normal cells are required for apoptosis induction of β-catCA- or GSK-3β-overexpressing cells. [Tg(hsp70l:GFP-T2A-β-catCA)] or [Tg(hsp70l:GFP-T2A-GSK-3β)] transgenic lines were exposed to heat shock. Percentages of embryos showing similar phenotype and number of embryos are shown. Scale bar, 200 μm. (G) Alleviation of Wnt/β-catenin activity difference between β-catCA-overexpressing cells and surrounding normal cells blocks β-catCA-overexpressing cell elimination. Cell membrane was visualized by injecting membrane-tagged mKO2 mRNA (red). Scale bar, 200 μm. (H) Cells causing excess noise in Wnt/β-catenin-gradients efficiently undergo apoptosis. Means ± SEM of caspase-3-active cell frequencies within the divided range along the AP axis (GSK-3βDN, n = 8 embryos, 1131 cells; Axin1, n = 14 embryos, 1308 cells) (bottom). See also Figure S3.

Zebrafish embryos consist of epithelial (envelope layer) and mesenchymal tissues (deep cells) (Carvalho and Heisenberg, 2010). In both tissues, Wnt/β-catenin-hyperactivated cells activated caspase-3 and then disappeared (Figures S3C and D), indicating global abnormal cell elimination. Mosaic but not ubiquitious introduction of abnormal Wnt/β-catenin activation or inhibition into normal embryos strongly activated caspase-3 (Figure 2F). Mosaically introduced β-catCA-expressing cells were eliminated from normal embryos but not from Wnt/β-catenin-hyperactivated embryos injected with antisense morpholino (MO) against the Wnt/β-catenin negative regulator APC or treated with the GSK-3β chemical inhibitor BIO (Figures 2G and S3D). Small numbers of β-catCA-expressing cells transplanted into normal but not β-catCA-expressing embryonic tissue activated caspase-3 in cells contacting normal cells (Figure S3E). These results indicate that Wnt/β-catenin activation or inhibition is not sufficient to induce apoptosis and suggest that Wnt/β-catenin-abnormal cells require surrounding healthy cells with appropriate Wnt/β-catenin activity for apoptosis induction. The Wnt/β-catenin activity difference between the abnormal and neighboring cells thus appears to trigger apoptosis in the former. Accordingly, cells with abnormally high or low Wnt/β-catenin activity efficiently activated caspase-3 in the Wnt/β-catenin signaling-low anterior or signaling-high posterior region, respectively (Figure 2H). This result indicates that a large Wnt/β-catenin activity differential between abnormal and neighboring cells is required for abnormal cell apoptosis induction. Mosaic activation of neither Ras nor Src signaling could activate caspase-3 in zebrafish embryos (Figure S3F), indicating that zebrafish embryonic tissue specifically kills Wnt/β-catenin-abnormal cells. However, survival of keratinocytes with abnormally high and low Wnt/β-catenin activity in zebrafish larval skin (Figures S3G and S3H) suggests that the Wnt/β-catenin-abnormal cell elimination system may specifically exist in tissues forming Wnt/β-catenin-gradients.

### Membrane β-Catenin and Cadherin Proteins Form Concentration Gradients Along the AP-Axis in a Wnt/β-catenin Gradient-Dependent Manner

Next, we investigated the mechanisms for sensing cells with unfit Wnt/β-catenin activity. Mosaic introduction of constitutive active or dominant negative β-catenin nuclear effector Lef1 (Lef1CA and Lef1DN) mutants did not activate caspase-3 (Figures 3A and 3B). Lef1DN co-expression did not block caspase-3 activation in β-catCA-expressing cells (Figures 3A and 3B), suggesting that unfit Wnt/β-catenin signaling triggers apoptosis independent of Lef1. Accordingly, β-catCA proteins localized to the nucleus, cytoplasm, and membrane (Figure 3C), whereas mosaic nuclear-export-signal (NES)-tagged β-catCA expression, which mainly localized to the cytoplasm and membrane (Figure 3C), activated caspase-3 (Figures 3A and 3B). These results suggest that unfit Wnt/β-catenin signaling in the cytoplasm or membrane, but not in the nucleus, triggers apoptosis.

**Figure 3.**
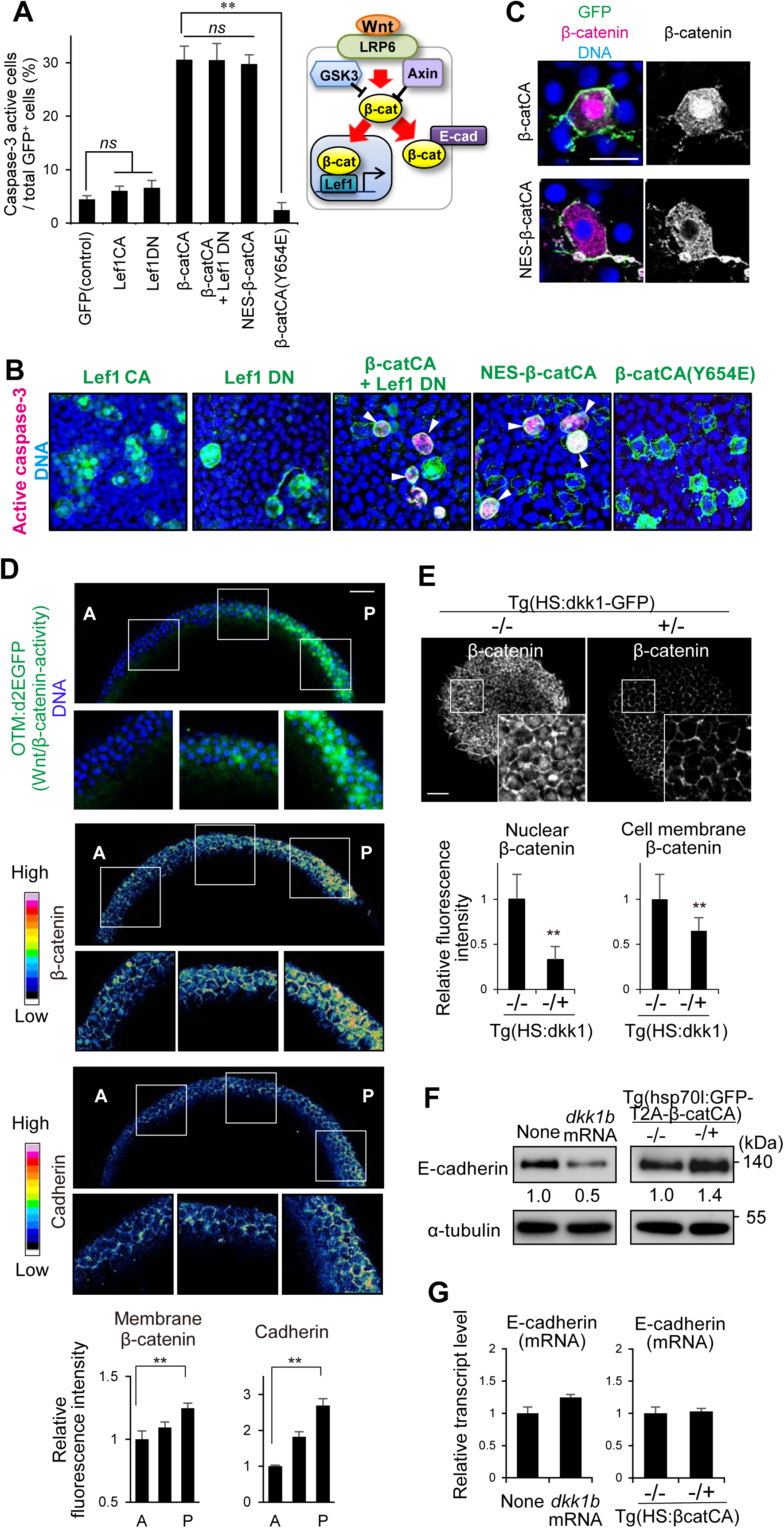
Membrane β-catenin and Cadherin Proteins Form their Concentration Gradient Along the AP-Axis in a Wnt/β-catenin Gradient-Dependent Manner. (A) Direct β-catenin-E-cadherin binding is required for unfit cell apoptosis induction. β-catenin transactivation with Lef1 and β-catenin cellular adhesion with E-cadherin (right). Means ± SEM of GFP^+^ caspase-3-active cell frequencies in Lef1 or β-catCA mutant mosaic embryos (left). \*\**p* < 0.01. (B) Representative confocal fluorescence images show GFP (green), active caspase-3 (magenta), and DNA (blue) in embryos introduced with cells expressing Lef1 mutants or β-catCA mutants. Scale bar, 50 μm. (C) Both β-catCA and NES-β-catCA localize to the plasma membrane in zebrafish embryos. Scale bar, 20 μm. (D) Membrane β-catenin and E-cadherin protein levels correlate with Wnt/β-catenin signaling activity. Optical sagittal cross-section (dorsal side) in 8.3 hpf embryos. Scale bar, 50 μm. Fluorescence intensity (means ± SEM) of β-catenin and E-cadherin staining at intercellular boundaries within 3 evenly divided regions across the AP axis. (Also see Methods). \*\**p* < 0.0l. (E) Inhibition of Wnt signaling (Dkk1 overexpression) reduces nuclear as well as membrane β-catenin. Dorsal side of whole-mount immunostaining of Tg(HS:dkk1b-GFP) zebrafish embryos and sibling embryos at 9 hpf exposed to heat shock at 37 °C from 4.3 to 5.3 hpf. Scale bar, 50 μm. Means ± SD. (F) E-cadherin protein level correlates with Wnt/β-catenin signaling activity. Embryo extracts at 9 hpf following heat shock at 37 °C from 4.3 to 5.3 hpf or *dkk1b* mRNA injection into one cell-stage embryos. (G) E-cadherin transcript level does not correlate with Wnt/β-catenin signaling activity in 9 hpf embryos by qRT-PCR.

Because β-catenin binds to the adhesion molecule cadherin in the membrane (Nagafuchi and Takeichi, 1989), we hypothesized that cadherin may facilitate unfit cell apoptosis. Mosaic introduction of mutant β-catCA Y654E, with transactivation but not cadherin-binding activity (Huber and Weis, 2001; Roura et al., 1999; van Veelen et al., 2011), could not activate caspase-3 (Figures 3A and 3B). This result suggests that direct β-catenin-cadherin interaction may be required to induce unfit cell apoptosis. Immunostaining of membrane β-catenin and cadherin expression in zebrafish embryos detected Wnt/β-catenin activity (OTM:d2EGFP) gradient formation and nuclear β-catenin level as well as membrane β-catenin and cadherin gradient formation along the AP axis (Figure 3D). Cells in the Wnt/β-catenin-high posterior region expressed high nuclear and membrane β-catenin and cadherin levels, albeit relatively low levels in the Wnt/β-catenin-low anterior region (Figure 3D), suggesting that Wnt/β-catenin signaling may promote membrane β-catenin and cadherin accumulation. Accordingly, Wnt antagonist Dkk1 overexpression in whole embryos reduced membrane β-catenin (Figure 3E) and E-cadherin protein levels (Figure 3F), whereas β-catenin overexpression increased E-cadherin protein levels (Figure 3F). Conversely, Dkk1 or β-catenin overexpression did not affect E-cadherin mRNA levels (Figure 3G), suggesting that Wnt/β-catenin signaling post-translationally stabilizes E-cadherin in zebrafish embryos.

### Cadherin is Involved in Unfit Cell Sensing

As expected, upon introducing abnormally high Wnt/β-catenin activity into the cadherin level-low anterior tissue, Wnt/β-catenin-abnormally-high cells increased both endogenous and exogenous cadherin proteins, with the converse also being true (Figures 4A, 4B, S4A–C). These results suggest that unfit cells alter cadherin levels post-translationally to yield substantial differences of membrane cadherin levels (cadherin imbalance) between unfit and neighboring normal cells. We hypothesized that this imbalance may be involved in unfit cell elimination. To test this, we relieved the cadherin imbalance by partial knockdown (Figure S4D) or E-cadherin overexpression in whole embryos. Both treatments reduced Wnt/β-catenin abnormally high- and low- cell apoptosis (Figures 4C and 4D). Mosaic introduction of cells overexpressing E-cadherin was sufficient to induce caspase-3 activation (Figure 4E). To avoid β-catenin absorption (and consequent Wnt/β-catenin activity reduction) by overexpressing E-cadherin, we also examined the E-cadherin αC mutant lacking β-catenin-binding activity. E-cadherin αC mutant-expressing cells also activated caspase-3 (Figure 4E), suggesting that abnormally high levels of cadherin stimulate apoptosis downstream of unfit β-catenin signaling. In addition, small numbers of E-cadherin-knockdown cells transplanted into normal embryonic tissue activated caspase-3 (Figure 4F), suggesting that abnormally low levels of cadherin also stimulate apoptosis. Thus, the membrane cadherin level difference between unfit and neighboring fit cells appears to be involved in unfit cell sensing.

**Figure 4.**
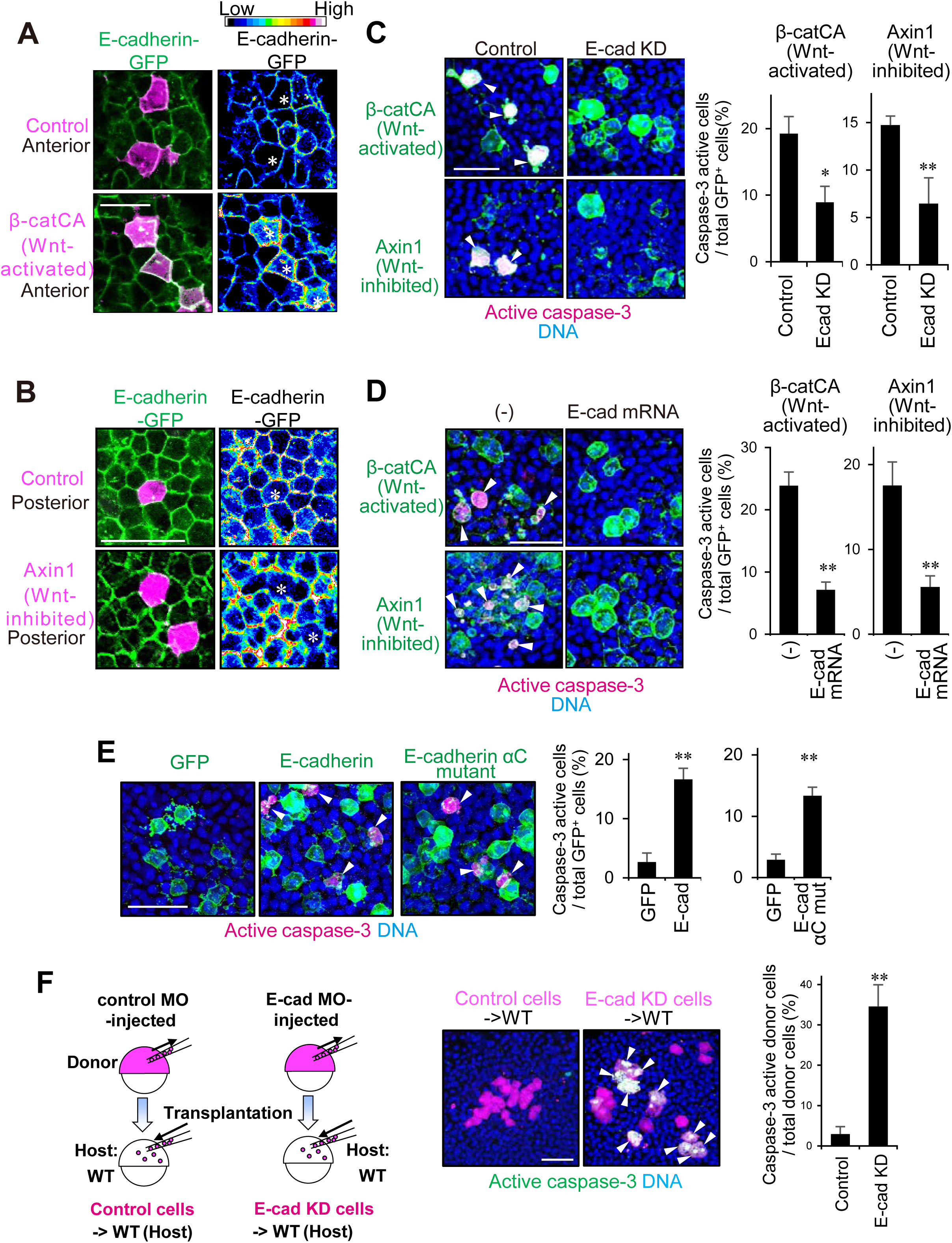
Cadherin is Involved in Unfit Cell Sensing. (A, B) Mosaic unfit abnormal Wnt/β-catenin-activity cell introduction [β-catCA- (Wnt-activated) (A) or Axin1- (Wnt-inhibited) (B) overexpressing] changes E-cadherin levels in zebrafish embryos. (C) Partial E-cadherin knockdown blocks β-catCA- or Axin1-overexpressing cell apoptosis. Scale bar, 50 μm. Means ± SEM of GFP^+^ (β-catCA, Axin1) caspase-3 active cell frequencies (right). \*\**p* < 0.01, \**p* < 0.05. (D) E-cadherin overexpression in zebrafish embryos blocks β-catCA- or Axin1-overexpressing cell apoptosis. Scale bar, 100 μm. Means ± SEM of GFP^+^ (β-catCA, Axin1) caspase-3 active cell frequencies. \*\**p*<0.01. (E) Cells with unfit high E-cadherin levels undergo apoptosis in mosaic embryos. Scale bar, 50 μm. Means ± SEM of GFP^+^ caspase-3-active cell frequencies (right). \*\**p* < 0.01, \**p* < 0.05. (F) Cells with unfit low E-cadherin levels undergo apoptosis in mosaic embryos. E-cadherin knockdown (E-cad KD) cells or control cells were transplanted into wild type embryos. Scale bar, 50 μm. Means ± SEM of GFP^+^ caspase-3-active cell frequencies (right). \*\**p* < 0.01. See also Figure S4.

### TGF-β-Smad Signaling Mediates Unfit Cell Killing

We next explored the unfit cell-killing system. As previous studies suggest a role for JNK MAPK, p38 MAPK, or p53 in cell competition-mediated unfit cell apoptosis (de la Cova et al., 2014; Moreno et al., 2002; Norman et al., 2012), we tested their potential involvement in Wnt/β-catenin-abnormal cell apoptosis in zebrafish embryos. β-catCA-expressing cells activated caspase-3 regardless of dominant negative JNK or p38 mutant expression (Figures S5A and S5B), treatment with chemical inhibitors against JNK and p38, or MO-mediated p53 knockdown (Figures S5C and S5D). Thus, JNK, p38, and p53 appear not to mediate unfit-Wnt/β-catenin-activity cell apoptosis.

To identify the actual mediators, we screened the genes involved in unfit cell-killing. We collected β-catCA-expressing cells from β-catCA-mosaically introduced (Mosaic) or β-catCA ubiquitously-expressing (Ubiquitous) embryos, or cells from uninjected embryos (Uninjected) (Figure 5A), and compared gene expression patterns using RNA-Seq (Figures 5A and S5E). Up- or downregulated genes in “Mosaic” but not in “Ubiquitous”, compared with “Uninjected” cells should be due to unfit cell introduction rather than simple Wnt/β-catenin activation. Notably, a negative regulator of Smad signaling, *skilb,* was downregulated in “Mosaic” cells, whereas several Smad-target genes (Lachmann et al., 2010) including *lnx1*, *foxo3b*, *sesn3*, *h2afy2*, *ccng1*, and*pkdccb*, were upregulated in unfit cells (Figures 5A and S5E, check-marks), suggesting upregulated Smad signaling in unfit cells. Smad family proteins consists of R-Smad (Smad1/2/3/5/8), co-Smad (Smad4), and I-Smad (Smad6/7). Upon stimulation with TGF-β superfamily of secreted ligands including TGF-βs and BMPs, R-Smad is phosphorylated and forms a complex with Smad4, which translocates into the nucleus to activate target gene expression. In general, Smad2/3 mediate TGF-β signaling, whereas Smad1/5/8 mediates BMP signaling (Gaarenstroom and Hill, 2014). A TGF-β-type Smad, Smad2, translocated into the nucleus in both Wnt/β-catenin abnormally high and low cells (Figure 5B). SBE-luc, which expresses luciferase in response to TGF-β-type Smads-dependent signaling pathway activation, was also activated in both abnormal cells (Figure 5C). In contrast, Smad2 nuclear translocation and SBE-luc activation was not detected in AP-axis forming zebrafish embryos (Figures 5B and 5C). Smad1/5/8 phosphorylation, which is activated in zebrafish presumptive ventral tissue (Figure S5F) to promote dorsoventral axis formation (Eivers et al., 2008), was not detected in Wnt/β-catenin unfit cells (Figure S5G). These results indicate that the unfit cells activate TGF-β-Smad signaling, which is not activated in normal embryonic tissue, but not BMP-Smad signaling, which controls embryonic dorsoventral patterning in parallel with Wnt/β-catenin signaling.

**Figure 5.**
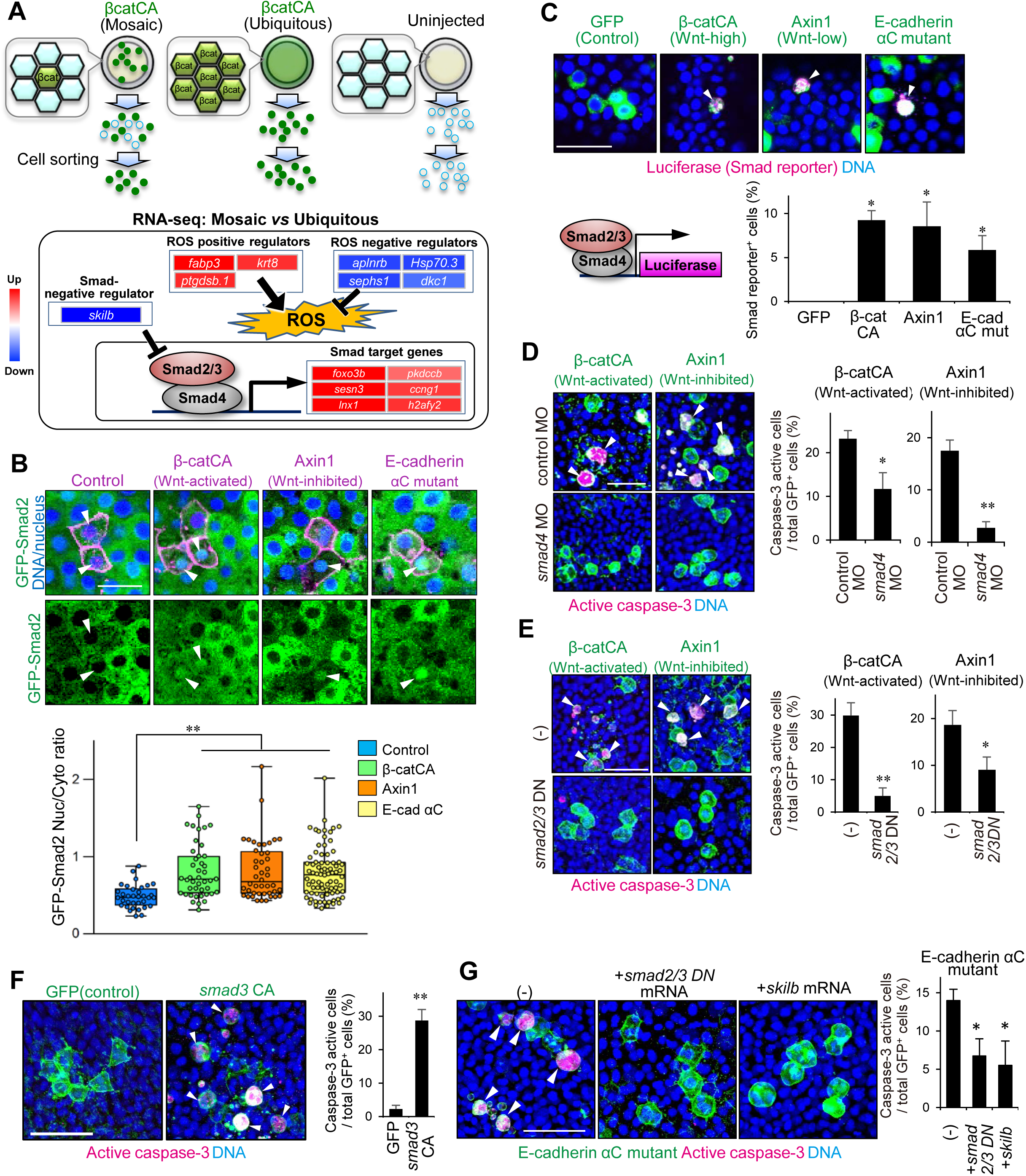
TGF-β-Smad Signaling Mediates Unfit Cell Killing. (A) Schematic representation of mRNA expression changes by RNA-Seq between β-catCA-mosaically introduced (Mosaic), ubiquitously β-catCA-expressing (Ubiquitous), or uninjected embryos. (See also Figure S6E). (B) GFP-Smad2 translocates into unfit Wnt/β-catenin-activity cell nuclei. Confocal fluorescence images of cells overexpressing indicated genes in GFP-Smad2 mRNA (200 pg)-injected embryos. Box plots of nuclear/cytoplasmic GFP-Smad2 ratio show 75th, 50th (median), and 25th percentiles (right). Whiskers indicate minimum and maximum. Each dot represents one embryo. \*\**p* < 0.01. (C) Wnt/β-catenin-abnormal cells activate a Smad2/3/4-dependent reporter gene. Embryos injected with heat shock-driven constructs. Scale bar, 50 μm. Smad reporter active cell frequencies are graphed. \**p* < 0.05. (D, E) Smad4 knockdown (D) or forced expression of *smad2/3* dominant negative mutants (*smad2/3*DN) (E) blocks βcatCA- or Axin1-expressing cell apoptosis. Means ± SEM of GFP^+^ (β-catCA- or Axin1-expressing) caspase-3-active cell frequencies (right). \*\**p* < 0.01, \**p* < 0.05. (F) Mosaic Smad3 activation is sufficient to induce apoptosis. Means ± SEM of GFP^+^ caspase-3-active cell frequencies (right). \*\**p* < 0.01. (G) E-cadherin αC mutant-expressing cell apoptosis is inhibited in *smad2/3*DN- or *skilb*-overexpressing embryos. Whole-mount immunostaining. Scale bar, 50 μm. Means ± SEM of GFP^+^ caspase-3-active cell frequencies (right). \**p* < 0.05. See also Figure S5.

We next examined TGF-β-Smad signaling involvement in unfit cell-killing. TGF-β-Smad signaling inhibition by *smad4* MO (Figure 5D), *smad2/3* DN mutant mRNA (Figure 5E), or *skilb* mRNA (Figure S5H) injection reduced both Wnt/β-catenin-abnormally-high and -low cell apoptosis. Mosaic introduction of cells expressing a constitutively active Smad3 mutant (Smad3CA) activated caspase-3 (Figure 5F). Mosaic introduction of the E-cadherin αC mutant activated Smad2 nuclear translocation (Figure 5B) and SBE-luc (Figure 5C) whereas *smad2/3* DN mutants or *skilb* overexpression blocked E-cadherin αC mutant-induced caspase-3 activation (Figure 5G). These results suggest that TGF-β-Smad signaling mediates unfit cell-killing downstream of cadherin.

### ROS Kills Unfit Cells Downstream of Cadherin and Smad Signaling

Smad signaling promotes ROS production (Liu and Desai, 2015). *foxo3b* and *sesn3*, Smad-target genes upregulated in unfit cells (Figures 5A and S5E), are also involved in ROS-mediated apoptosis (Hagenbuchner et al., 2012). RNA-Seq data also showed that negative regulators of ROS (*aplnrb*, *hsp70.3*, *sephs1*, and *dkc1*) (Gu et al., 2011; Shi et al., 2016; Than et al., 2014; Tobe et al., 2016) were downregulated in unfit cells, whereas positive regulators (*fabp3*, *ptgdsb.1*, and *krt8*) (Mathew et al., 2008; Shen et al., 2012; Sugita et al., 2013) were upregulated (Figures 5A and S5E). Sephs1 expression was downregulated in both Wnt/β-catenin abnormally-high and -low cells, as determined by immunostaining (Figure 6A). ROS may thus be activated in unfit cells. Accordingly, mosaic but not ubiquitous introduction of abnormal Wnt/β-catenin activation or inhibition into normal embryos activated ROS production (Figure 6B, Figure S6). Co-expression of the ROS negative regulators SOD1 (Greenlund et al., 1995) or Sephs1 (Tobe et al., 2016), or *smad4* knockdown reduced ROS levels in β-catCA-expressing cells (Figure 6B). ROS were also produced in cells expressing Smad3CA or the E-cadherin αC mutant (Figure 6B). These results suggest that cells with unfit Wnt/β-catenin activity activate ROS production through cadherin and Smad signaling. Furthermore, SOD1 or Sephs1 overexpression blocked caspase-3 activation in the artificially introduced Wnt/β-catenin abnormally-high and -low cells (Figure 6C), suggesting that ROS mediates unfit cell apoptosis.

**Figure 6.**
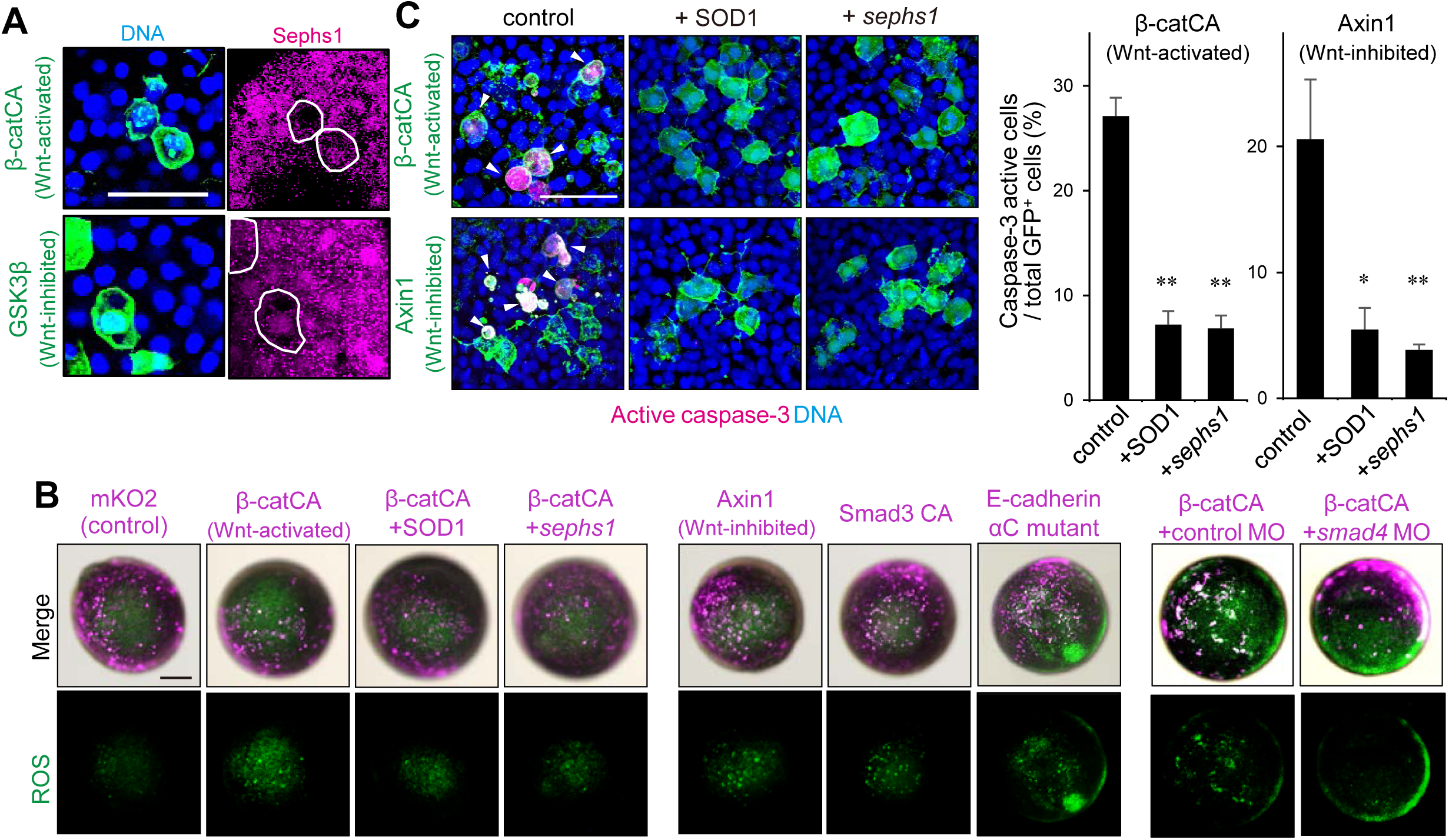
ROS Kills Unfit Cells Downstream of Cadherin and Smad Signaling. (A) Representative confocal fluorescent images show GFP (green) with β-catCA or GSK3β, Sepsh1 (magenta) and DNA (blue). (B) Cells with unfit Wnt/β-catenin activity activate ROS production through E-cadherin and Smad signaling. Fluorescence images showing ROS probe (CellRox Green)-stained mosaic embryos. Scale bars, 200 μm. (C) ROS downregulation blocks β-catCA- or Axin1-expressing (Wnt-activated or -inhibited) cell apoptosis. Confocal fluorescence images in mosaic embryos. Scale bar, 200 μm (left). Means ± SEM of GFP+ caspase-3-active cell frequencies (right). \*\**p* < 0.01, \**p* < 0.05. See also Figure S6.

### Apoptotic Elimination of Unfit Cells is Required for Precise Tissue Patterning

To confirm the importance of Wnt/β-catenin-abnormal cell elimination for embryogenesis, we examined the effects of unfit cell apoptosis inhibition. Blocking apoptosis by p35 co-expression in cells with artificially introduced β-catCA induced ectopic expression and abnormal reduction of brain AP markers (Figure S7A). Co-expression of p35 with β-catCA or GSK-3β increased abnormal morphogenesis (Figure S7B), whereas expression of p35, β-catCA, or GSK-3β alone had little effect.

To further test the physiological significance of unfit cell elimination, we blocked ROS-mediated unfit cell apoptosis by overexpressing a ROS negative regulator. Sephs1 overexpression partially reduced physiologically occurring apoptosis (Figure 7A). SOD1 or Sephs1 overexpression induced both ectopic activation and abnormal reduction of the Wnt/β-catenin reporter (Figure 7B) and brain AP markers (Figure 7C) and disturbed embryonic morphogenesis (Figure 7D), as did treatment with the ROS scavenger *N*-acetyl-L-cysteine (NAC) (Figure 7B). These data suggest that ROS-mediated elimination of naturally generated Wnt/β-catenin-unfit cells is essential to achieve smooth Wnt/β-catenin-gradient formation and precise embryogenesis.

**Figure 7.**
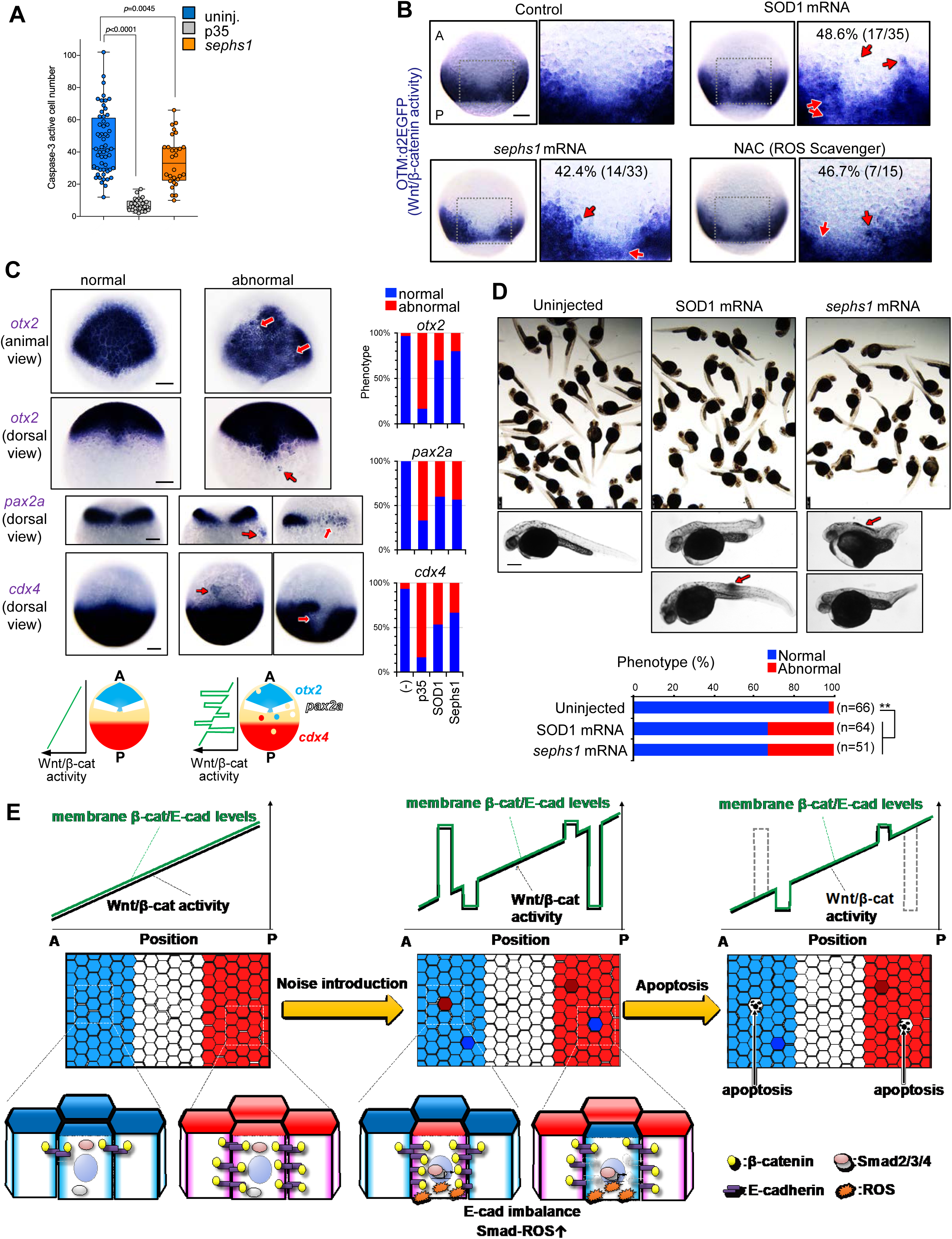
Apoptotic Elimination of Unfit Cells is Required for Precise Tissue Patterning. (A) Sephs1 overexpression partially reduces physiologically occurring apoptosis in embryos as determined by caspase-3 immunostaining. Box plots of caspase-3-active cell number per embryo at 10 hpf stage show 75th, 50th (median), and 25th percentiles. Whiskers indicate minimum and maximum. Each dot represents one embryo. (B) Inhibiting ROS increase distorts the Wnt/β-catenin-gradient. Whole-mount *in situ* hybridization of d2EGFP in Tg(OTM:d2EGFP) embryos (dorsal view) injected with *SOD1* or *sephs1* mRNA (800 pg) or treated with 100 μm NAC. Magnification of boxed area (black line) (right). Embryo percentages and numbers with similar expression patterns are shown. Red arrows: ectopic activation or inactivation areas. Scale bar, 200 μm. (C) Inhibition of ROS-mediated apoptosis distorts AP patterning. Whole-mount *in situ* hybridization of *otx2* (marker of presumptive forebrain and midbrain), *pax2a* (marker of presumptive midbrain-hindbrain boundary), and *cdx4* (marker of presumptive spinal cord) in embryos uninjected or injected with *p35*, *sephs1*, or *SOD1* mRNA (800 pg). Schematic illustration indicates expression pattern of AP tissue markers. Percentages of embryos with normal or abnormal expression patterns are shown. Scale bar, 200 μm. (D) Overexpression of *SOD1* or *sephs1* mRNA induces abnormal morphogenesis. Images show 32 hpf zebrafish larvae uninjected or injected with *SOD1* or *sephs1* mRNA (800 pg). Red arrow indicates abnormal cell proliferation. Scale bar, 500 μm. Percentages of embryos with normal or abnormal morphology are shown. The numbers shown above the graph indicate the total numbers of embryos analyzed. \*\**p* < 0.01. (E) Schematic diagram of the Wnt/β-catenin-noise canceling system. See also Figure S7.

## DISCUSSION

The present study demonstrates a cell competition-mediated “noise-canceling system” for morphogen signaling-gradients (Figure 7E). During zebrafish embryonic AP patterning, membrane β-catenin and cadherin protein level gradients along the AP axis are formed in a Wnt/β-catenin activity-dependent manner. The spontaneous appearance of cells with unfit Wnt/β-catenin activity produces substantial noise in the Wnt/β-catenin-gradient and alters membrane β-catenin and cadherin protein levels, leading to cadherin level imbalance between unfit and neighboring cells. Unfit cells also activate TGF-β-Smad signaling to produce ROS and consequently undergo ROS-mediated apoptosis. In this manner, embryonic tissues eliminate excess noise from the Wnt/β-catenin gradient to support its proper formation along with robust embryonic patterning.

Cells artificially introduced with abnormally high or low BMP signaling activity undergo apoptosis through cell competition or its related system in *Drosophila* imaginal tissues and mouse epiblasts (Adachi-Yamada et al., 1999; Moreno et al., 2002; Sancho et al., 2013). However, the physiological importance and detailed mechanisms of this phenomenon remain unclear. We show that both artificially and physiologically introduced unfit cells, which produce excess noise in the Wnt/β-catenin morphogen-gradient, undergo apoptosis through cadherin, TGF-β-Smad, and ROS, and reveal the physiological importance of this system in signaling-gradient formation and embryonic morphogenesis. Thus, we unraveled an undescribed morphogen-gradient-correcting system that plays physiological roles in normal development. The previous findings together with our present results suggest that the apoptosis-mediated morphogen noise-canceling system may be evolutionarily conserved, with Wnt/β-catenin and other morphogen signaling utilizing similar systems. ROS signaling inhibition partially reduced the number of physiologically appearing apoptotic cells, suggesting that these cells comprise Wnt/β-catenin-defective (ROS-activated) cells along with other cells, potentially including other morphogen signaling-defective cells. With regard to sensing “unfitness”, only one report has identified that a ligand protein Sas and its receptor PTP10D are involved in recognizing polarity-deficient cells (Yamamoto et al., 2017). We showed that unfit Wnt/β-catenin activity is translated to unfit cadherin levels, which are sensed by neighboring cells with fit cadherin levels and fit Wnt/β-catenin activity. Apoptosis-independent morphogen gradient-correcting systems also exist. In the zebrafish developing neural tube in which Shh signaling forms its activity-gradient along the dorsal-ventral axis, cells with unfit Shh signaling activity migrate to the appropriate area to form a smooth Shh signaling-gradient (Xiong et al., 2013). Morphogen signaling pathways equip negative feedback systems mediated through negative signaling regulators such as Axin2 (in Wnt/β-catenin signaling), Smad6/7 (in BMP signaling) and patched (in Shh signaling) ( Ashe and Briscoe, 2006; Li et al., 2018). Such feedback systems also contribute to correcting morphogen signaling noise. Developing tissues may use these correction systems differently in accordance to the “unfitness” of unfit cells. Because the apoptosis-mediated system is activated only in cells causing substantial noise, this system might contribute in particular to eliminating cells with severe signaling-defects (e.g., with mutations that constitutively activate or inhibit morphogen signaling).

The apoptosis-mediated Wnt/β-catenin-noise canceling system appears to comprise a kind of cell competition supporting precise tissue morphogenesis. Myc-high cells compete with and kill Myc-low cells in developing mouse embryos, supporting proper development (Clavería et al., 2013; Sancho et al., 2013). We also observed that the mosaic introduction of cells expressing Myc at high levels killed surrounding normal cells in zebrafish embryos (Figure S4E). However, this competition was cadherin- and Smad4-independent (Figures S4E and S5I), indicating that the Myc-mediated competition and Wnt/β-catenin-noise canceling system mechanisms differ. Although JNK and p38 regulate apoptosis induction in many cell competition systems (Igaki, 2009; Norman et al., 2012), they appear not to be required for the Wnt/β-catenin-noise canceling system, which is instead mediated by cadherin-imbalance and the Smad-ROS axis. Cadherin functions in both mechanical cell-cell adhesion and cell recognition (Halbleib and Nelson, 2006; Takeichi, 1991). Cells expressing low and high cadherin levels separate out when mixed *in vitro*, indicating that cells can recognize quantitative differences in cadherin levels (Steinberg and Takeichi, 1994). However, it remains unclear how zebrafish embryonic cells translate quantitative differences in cadherin levels (cadherin-imbalance) to “unfitness” and consequently determine to kill unfit cells. Because cadherin-mediated adhesion maintains tissue integrity ( Halbleib and Nelson, 2006; Takeichi, 1991), local cadherin level change in unfit cells would cause local loss of tissue integrity, which may induce mechanical stress in unfit cells, in turn stimulating the unfit cell-killing system. Accordingly, mechanical stress can activate TGF-β-type Smad3 (Furumatsu et al., 2013; Huang et al., 2014; Kunnen et al., 2017; Saha et al., 2008), which act downstream of cadherin in the Wnt/β-catenin-noise canceling system. However, the detailed mechanisms by which cadherin-imbalance activates TGF-β-type Smad3 require further investigation.

The Wnt/β-catenin-noise canceling system may act context-dependently because cell clones with abnormally high Wnt/β-catenin activity can kill surrounding normal cells in *Drosophila* epithelial tissue (wing disc) (Vincent et al., 2011), whereas Wnt/β-catenin activity-aberrant keratinocytes survive and persist in zebrafish larval skin. Clone size, cell/tissue-type, and/or intra-tissue localization might affect unfit cell behavior. We consider that the Wnt/β-catenin-noise canceling system acts in proliferative tissues that form a Wnt/β-catenin-gradient and kills suddenly introduced single unfit cells that produce excess noise in the gradient. Therefore, this system might also function in the intestinal crypt, which undergoes active cell turnover and also forms a Wnt/β-catenin-gradient (Farin et al., 2016). Accordingly, in mouse intestine, mosaically introduced Wnt signaling-hyperactivated cells strongly express E-cadherin and actively undergo apoptosis (Wong et al., 1998), indicating that the mammalian intestine may possess a similar system.

Cells with abnormally high Wnt signaling may represent an origin of cancer (Kinzler et al., 1991; Nishisho et al., 1991). The Wnt/β-catenin-noise canceling system may therefore also eliminate Wnt/β-catenin-hyperactivated precancerous cells to prevent primary tumorigenesis. E-cadherin and Smad2/4, the mediators of this system, also act as tumor suppressors ( Christofori and Semb, 1999; Derynck et al., 2001). Their loss of function might reduce Wnt/β-catenin-noise canceling system activity, allowing the surviving Wnt/β-catenin-high precancerous cells to prime tumorigenesis. Thus, the Wnt/β-catenin-noise canceling system represents a facet of the roles of tumor suppressor genes. In the future, it would therefore be of interest to examine the roles of this system in cancer prevention.

## SUPPLEMENTAL INFORMATION

Supplemental information includes – Figures, -- Tables, and – Movies and can be found with this article online at:

## ACKNOWLEDGMENTS

We thank A. Kikuchi, E. Raz, A. Nagafuchi, M. Hibi, G. Salvesen, E. Fisher, W. El-Deiry, Y. Fujita, A. Yoshimura, S. Korsmeyer, M. Okada, and K. Kawakami for providing the plasmids; ZIRC and NBRP for providing transgenic zebrafish; A. Nagafuchi for helpful discussion; the Advanced Computational Scientific Program of the Research Institute for Information Technology, Kyushu University, Y. Sato in Nagoya University Live Imaging Center, Y. Kamei and T. Yabe in NIBB, J. Konno and Y. Kamihara in Olympus, NBRP zebrafish, and Ishitani lab members (Y. Sado, H. Okumura, M. Matsuo, M., Sakuma, K., Taniguchi, Y. Haraoka, M. Oginuma, and C. Mogi) for their technical support and fish maintenance. This research was supported by the Sumitomo Foundation and Takeda Foundation (T.I.), JSPS Research Fellowships for Young Scientists (Y.A.), JST CREST (JPMJCR16G1) (Y. O.), and a Grant-in-Aid for Scientific Research on Innovative Areas (25117720) (T.I.) (25116010) (Y.O.) and Scientific Research (B) (25293072) (T.I.).

## AUTHOR CONTRIBUTIONS

Conception and design: Y.A. and T.I.; Investigation: Y. A., S. O., H. F., S. I., R. A., J. N., T. M., N. S., Y. O., and T. I.; Bioinformatics and data analysis: J.N. and Y.O.; Writing and review: Y.A. and T.I.; Writing contribution and review: S. O., H. F., S. I., R. A., J. N., T. M., N. S., and Y. O.

## DECLARATION OF INTERESTS

The authors declare no competing interests.

## METHODS

### CONTACT FOR REAGENT AND RESOURCE SHARING

Further information and requests for resources and reagents should be directed to and will be fulfilled by the Lead Contact, Tohru Ishitani.

### EXPERIMENTAL MODEL AND SUBJECT DETAILS

#### Zebrafish maintenance

Zebrafish were raised and maintained under standard conditions. Wild-type strains (AB and India) were used, and the following transgenic lines were used: Tg(OTM:d2EGFP) (Shimizu et al., 2012), Tg(OTM:Eluc-CP), Tg(hsp70l:GFP-T2A-β-catCA), Tg(hsp70l:GFP-T2A-GSK-3β), and Tg(HS:dkk1b-GFP) (Stoick-Cooper et al., 2007). All experimental animal care was performed in accordance with institutional and national guidelines and regulations. The study protocol was approved by the Institutional Animal Care and Use Committee of the respective universities (Kyushu University Permit# A28-005-1; Gunma University Permit# 17-051). One-cell stage embryos were used for cell injection to generate transgenic fish or mosaic embryos, with the latter processed up to 9 hpf. One- and two cell-stage embryos were used for MO injection. Alternately, cells were injected into 3.3 hpf to 3.7 hpf stage embryos.

### METHOD DETAILS

#### Mosaic Wnt/β-catenin-abnormal cell introduction

Hsp70 promoter-driven plasmids (5–17.5 pg) were injected into one-cell-stage embryos and maintained at 28.5 °C until 4.3 hpf (dome stage). At 4.3 hpf, embryos were exposed to heat-shock. Briefly, embryos were transferred to pre-warmed egg water at 37 °C and kept at 37 °C for 1 hour. After heat-shock, embryos were placed at 28.5 °C, then fixed at 9 hpf for immunostaining or *in situ* hybridization.

#### Embryonic cell preparation for RNA-Seq

β-catCA-mosaically introduced embryos were prepared as described above. HS:GFP-2A-β-catCA-transgenic zebrafish embryos were used as β-catCA-ubiquitously introduced embryos. Cell dissociation was performed as previously described (Link et al., 2006) with the following modification: 40 embryos per group at 8.3 hpf stage were placed in a solution of 2 mg/ml Pronase (Roche, Upper Bavaria, Germany) in E2 (15 mM NaCl, 0.5 mM KCl, 2.7 mM CaCl_2_, 1 mM MgSO_4_, 0.7 mM NaHCO_3_, 0.15 mM KH_2_PO_4_, and 0.05 mM Na_2_HPO_4_) on a 2% agar-coated dish for approximately 6 min at 28.5 °C. After dechorionation, embryos were washed with deyolking buffer (1/2 Ginzburg Fish Ringer without calcium: 55 mM NaCl, 1.8 mM KCl, and 1.25 mM NaHCO_3_). Embryos were transferred into a 1.5 ml tube and then the yolk was disrupted by pipetting with a 1000 μl tip. The embryos were shaken for 5 min at 1100 rpm to dissolve the yolk (Thermomixer, Eppendorf, Hamburg, Germany). Cells were pelleted at 300 g for 1 min and the supernatant discarded. Cell pellets were resuspended in FACSmax Cell Dissociation Solution (Genlantis, San Diego, CA). β-catCA-expressing (GFP+) cells were sorted by FACSAriaII (BD, Franklin Lakes, NJ). Uninjected negative control cells were also sorted under similar conditions. Sorted cells were pelleted and dissolved in TRIzol reagent (Invitrogen, Waltham, MA).

#### Generation of transgenic zebrafish

Plasmid DNA (OTM:Eluc-CP, hsp70l:GFP-T2A-β-catCA, or hsp70l:GFP-T2A-GSK-3β) along with Tol2 transposase mRNA were co-injected into one-cell stage wild-type zebrafish (AB) embryos. A transgenic fish was outcrossed with a wild-type fish to produce the founder line. To generate a transgenic line carrying a single transgene, a carrier was outcrossed with wild-type fish. Reporter transgenic fish were maintained as homozygous transgenic fish.

#### Time-lapse imaging and data analysis

For time-lapse luminescence live imaging, Tg(OTM:Eluc-CP) embryos were manually dechorionated by forceps and then mounted in 1% low melting agarose with 3 mM D-luciferin potassium salt (Wako) onto glass bottom dishes (Matsunami Glass, Osaka, Japan). Images were collected using an LV200 bioluminescence microscope (Olympus, Tokyo, Japan) equipped with a 20× 0.75 NA objective at 28 °C. Collected image data were processed by TiLIA2 software (Olympus) for removing signals of cosmic rays, and then analyzed using ImageJ with plug-ins developed by Olympus.

For time-lapse confocal live imaging, embryos were manually dechorionated by forceps and mounted in 1% low melting agarose with egg water onto glass bottom dishes. Live imaging was performed using an LSM700 confocal laser scanning microscope (Zeiss, Oberkochen, Germany) equipped with a 20× 0.8 NA objective. Two laser lines, 488 nm and 559 nm were used. The recording interval was 1 to 2 min. At each time point, 12 confocal slices through the z-axis were acquired. Collected image data were processed using Imaris software (Bitplane, Zurich, Switzerland).

#### Detection of ROS

To detect ROS production, zebrafish embryos were incubated with 1.67 μm CellRox Green (Thermo Fisher) in egg water for 30 min. After loading, embryos were washed three times in egg water and fixed with 4% paraformaldehyde.

#### NAC treatment

OTM:d2EGFP-transgenic zebrafish embryos were treated with 100 μm *N*-acetyl-L-cysteine (NAC) (Sigma) from the 75% epiboly to 85% epiboly stage and then fixed.

### QUANTIFICATION AND STATISTICAL ANALYSIS

#### Immunofluorescence

For the quantification of membrane β-catenin and E-cadherin levels, the fluorescence intensity in two intercellular areas per cell (n = 6 cells) in each region was measured. Analyzes were performed in duplicate.

#### Statistical analysis

Differences between groups were examined using a two-tailed unpaired Student *t* test, a one-way ANOVA test, and Fisher’s exact test in Excel (Microsoft, Redmond, WA) or Prism7 (GraphPad Software, San Diego, CA). *p* values <0.05 were considered significant. Details of statistical analysis and sample numbers are shown in Table S1.

---

**Figure S1.**
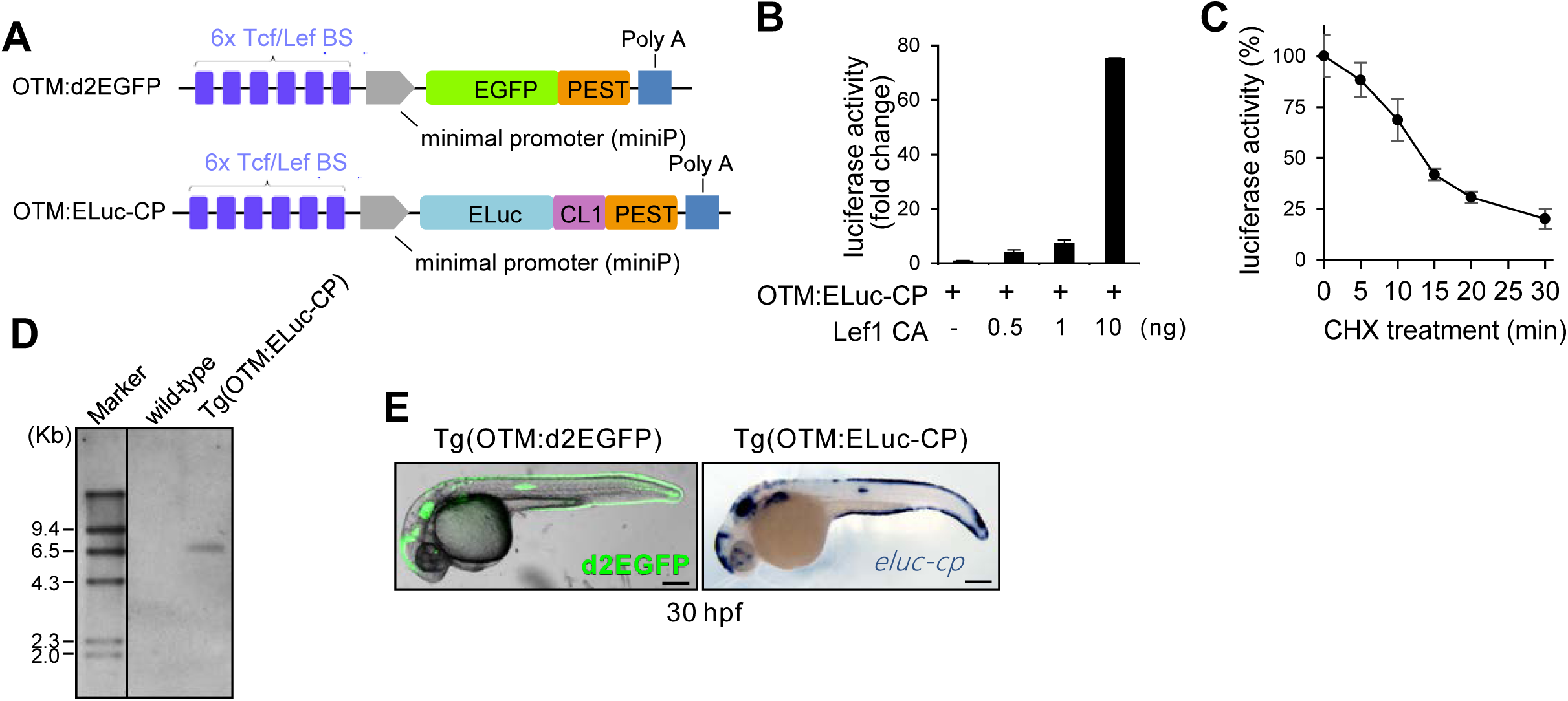
Wnt/β-catenin signaling reporter properties. Related to Figure 1. (A) Schematic diagrams of two Wnt/β-catenin signaling-reporter constructs: OTM:d2EGFP and OTM:ELuc-CP. Tcf/Lef BS: consensus sequence of the Tcf/Lef-binding site; EGFP: enhanced green fluorescent protein; ELuc: Emerald Luciferase; PEST: degradation sequence derived from mouse ornithine decarboxylase; CL1: degradation sequence derived from yeast; Poly A: SV40 polyadenylation sequence. (B) OTM:ELuc-CP drives ELuc-CP expression in mouse neuro-2a cells in response to the activation of Wnt/β-catenin signaling. Neuro-2a cells were transfected using a reporter construct with a titrated amount of an expression plasmid-encoding Lef1 CA. Then, luciferase activity was measured. (C) The half-life of ELuc-CP is approximately 15 min. Neuro-2a cells were transfected using a reporter construct. After 36 hours, cells were treated with 100 ng/ml cycloheximide (CHX) for the indicated times. Then, luciferase activity was measured. The error bars indicate the standard deviations. (D) The OTM:ELuc-CP-transgenic zebrafish line carries a single copy of the OTM:ELuc-CP reporter. Southern blot analysis of the transgenes in the OTM:ELuc-CP transgenic zebrafish line. Genomic DNA was prepared from the tail fins of this line and used for Southern blot analysis. (E) OTM:ELuc-CP and OTM:d2EGFP are activated in same areas of zebrafish embryos. Left panel shows d2EGFP fluorescence (green) merged with bright-field of OTM:d2EGFP-transgenic zebrafish at 30 hpf (left view with the anterior side to the left). Right panel shows whole mount *in situ* hybridization of *eluc-cp* (blue) in OTM:ELuc-CP-transgenic zebrafish at 30 hpf (left view with the anterior side to the left). Scale bar, 200 μm.

**Figure S2.**
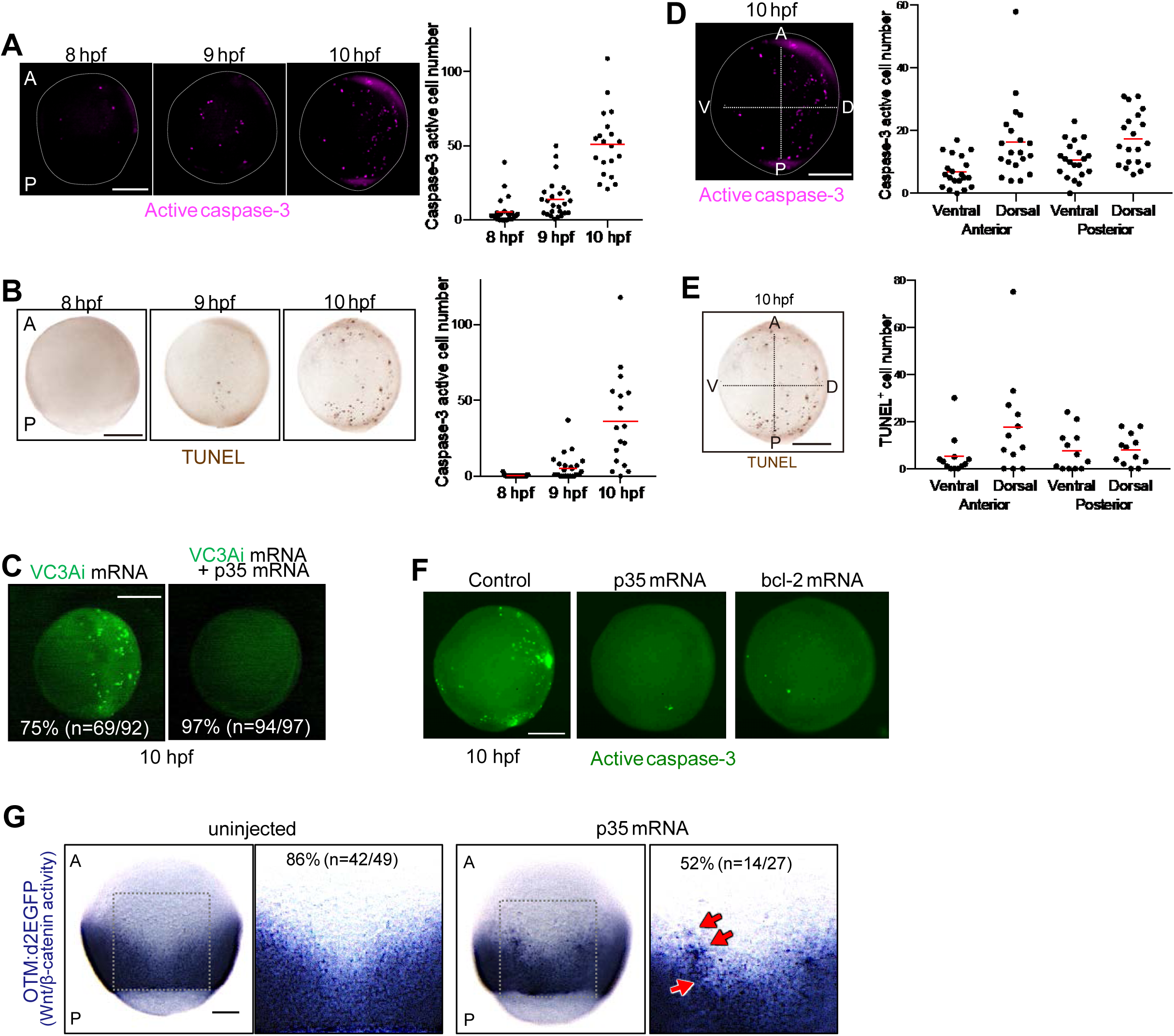
Physiological apoptosis during zebrafish AP axis formation. Related to Figure 1. (A) Caspase-3 activation during zebrafish AP axis formation. Representative fluorescence images showing whole-mount immunostaining of active caspase-3 (magenta) in embryos maintained with normal conditions. White dotted line indicates outline of the embryo (lateral view with the anterior to the top, ventral to the left). Numbers of caspase-3-active cells in each embryo are graphed. Each dot represents one embryo (8 hpf, n = 27 embryos; 9 hpf, n = 25; 10 hpf, n = 20). Means are indicated by a horizontal bar. Scale bar, 200 μm. (B) Cell death during zebrafish AP axis formation. Same analysis as in (A) with TUNEL assay (8 hpf, n = 29 embryos; 9 hpf, n = 27; 10 hpf, n = 17). (C) Live imaging of caspase-3 activation in zebrafish embryos. Representative fluorescence images show an embryo injected with mRNA of the caspase activity fluorescent biosensor/VC3Ai (Zhang et al., 2013) (250 pg) (green), or co-injected with the caspase inhibitor p35 (800 pg) (lateral with the anterior to the top, ventral to the left). Co-injection of p35 mRNA effectively reduced VC3Ai fluorescence, suggesting that the fluorescence in VC3Ai-injected embryos shows the activation of caspase. Scale bar, 200 μm. (See also Movie S2). (D) Caspase-3 active cells appear in an irregular manner. Representative fluorescence images showing whole-mount immunostaining of active caspase-3 (magenta) in embryos maintained with normal conditions at the 10 hpf stage (lateral view with the anterior to the top, ventral to the left). A, P, V, and D indicate anterior, posterior, ventral, and dorsal, respectively. Scale bar, 200 μm. The numbers of caspase-3-active cells in each embryo are graphed. Each dot represents one embryo (n = 20 embryos). The number of caspase-3-active cells/per embryos and the position of these cells were irregular. (E) Same analysis as in (D) with TUNEL assay (n = 17 embryos). (F) Overexpression of *bcl-2* or *p35* efficiently reduced physiologically occurring apoptosis. Representative fluorescence images showing whole-mount immunostaining of active caspase-3 (green) in embryos uninjected or injected with *bcl-2* or *p35* mRNA. Scale bar, 200 μm. (G) Apoptosis inhibition distorts the Wnt/β-catenin-gradient. Whole-mount *in situ* hybridization of d2EGFP in Tg(OTM:d2EGFP) embryos (dorsal view) uninjected or injected with p35 mRNA (800 ng). Magnified image of boxed area (black line; right). Embryo percentages and numbers with similar expression patterns are shown. Red arrows: ectopic activation or inactivation areas. Scale bar, 200 μm.

**Figure S3.**
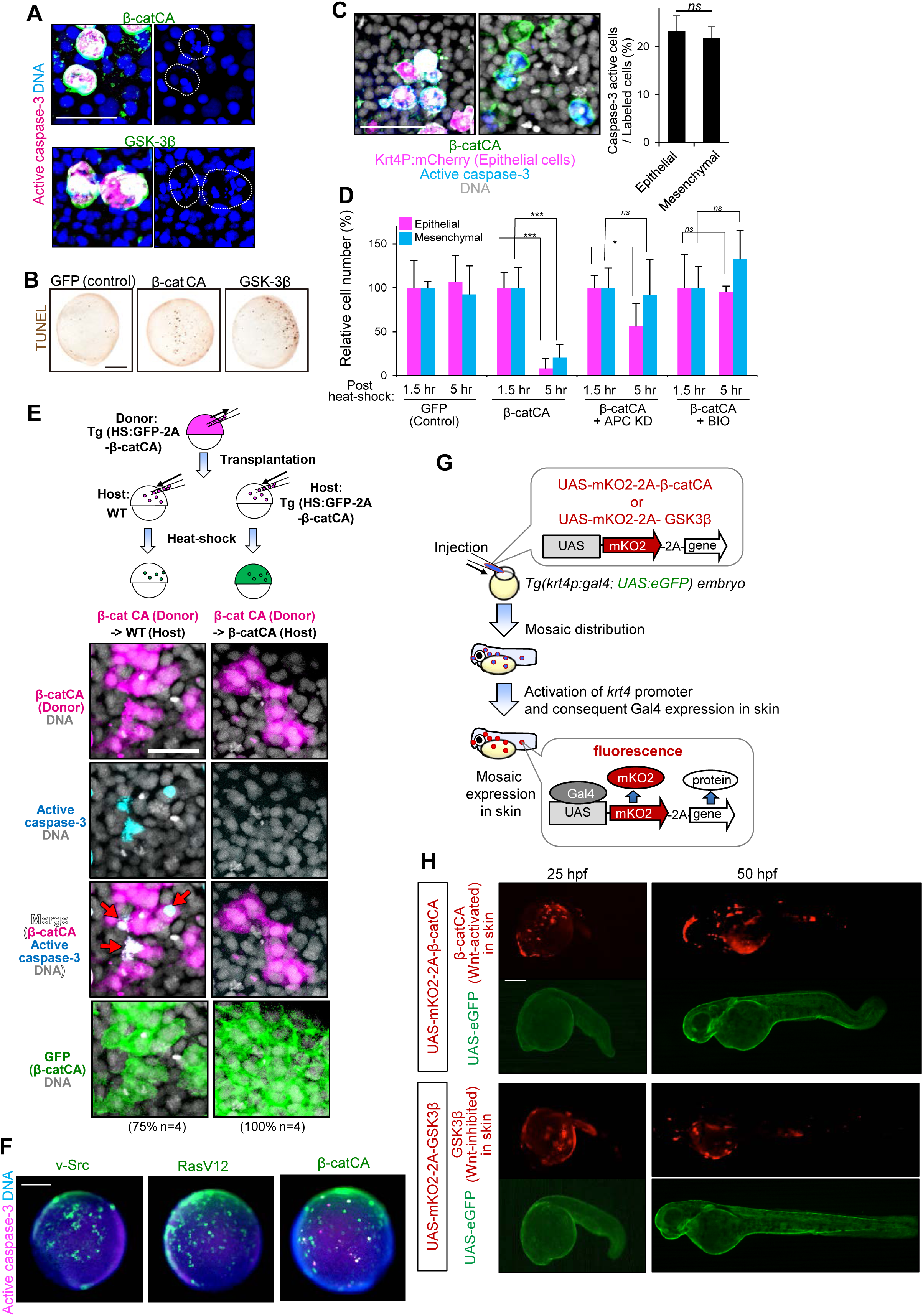
Wnt/β-catenin activity-abnormal cells require their neighboring normal cells to undergo apoptosis. Related to Figure 3. (A) DNA fragmentation occurs in caspase-3-active β-catCA-overexpressing cells or GSK-3β-overexpressing cells. Representative confocal fluorescence images show GFP (green), active caspase-3 (magenta), and DNA (blue) in embryos. Scale bar, 20 μm. (B) DNA fragmentation confirmed by TUNEL assay. Scale bar, 200 μm. (C) Apoptosis of β-catCA-overexpressing cells (cells with abnormally high Wnt activity) occurs both in epithelial cells (GFP+mCherry+) and mesenchymal cells (GFP^+^mCherry^−^). Embryos were co-injected with an Hsp70 promoter-driven β-catCA-expression plasmid (hsp70l:GFP-T2A-β-catCA) and Krt4P:mCherry plasmid, which expresses mCherry in epithelia. mCherry+ cells locate in the epithelia, whereas mCherry^−^ cells are mesenchymal cells. Means ± SEM of GFP^+^ caspase-3 active cell frequencies are shown. Scale bar, 50 μm. (D) β-catCA-overexpressing cells are eliminated from embryos, whereas upregulation of Wnt/β-catenin activity in whole embryos blocks cell elimination. Percentages of relative GFP+ cell numbers at 1.5 to 5 hours post heat-shock in APC knocked-down (KD) or 10 μm BIO-treated embryos are shown. \*\*\**p* < 0.001, \**p* < 0.05. (E) Transplantation of a small number of β-catCA-expressing cells into normal embryonic tissue activates caspase-3 in the transplanted cells in contact with normal cells. Schematic illustration of the experimental design for cell transplantation. Rhodamine-dextran-labeled donor cells from Tg(hsp70l:GFP-T2A-β-catCA) embryos were transplanted into unlabeled wild-type or Tg(hsp70l:GFP-T2A-β-catCA) host embryos. Representative confocal fluorescence images show donor cells (magenta), DNA (grey), active caspase-3 (cyan), and GFP/transgene carrier (green) in transplanted embryos. The percentages of embryos showing similar phenotype and number of embryos are shown under each image. Scale bar, 50 μm. (F) Signaling-abnormal cells expressing v-Src or RasV12 do not undergo apoptosis. Representative fluorescence images show whole mount immunostaining of GFP (green), active caspase-3 (magenta), and DNA (blue) in embryos mosaically introduced with cells expressing v-Src or RasV12. Scale bar, 200 μm. (G, H) Schematic illustration of experimental introduction of Wnt/β-catenin-abnormal cells in zebrafish larval skin. (G) Plasmids that express both membrane-tagged mKO2 and a Wnt/β-catenin-regulator gene in response to Gal4-mediated UAS promoter activation were injected into one-cell stage *Tg(krt4p:gal4; UAS:eGFP*) zebrafish embryos (gifts from Drs. K. Kawakami and H. Wada) (Asakawa et al., 2008). Cell division during development promotes mosaic distribution of injected plasmids. *Keratin4* (*Krt4*) promoter activates Gal4 expression in larval skin (keratinocytes). Consequently, keratinocytes with the plasmid express both mKO2 and the Wnt modulator gene, resulting in the appearance of a small number of mKO2^+^ Wnt/β-catenin-abnormal cells. *UAS:eGFP* is also activated in larval skin. (H) Fluorescence images show embryos artificially introduced with keratinocytes overexpressing β-catCA (Wnt/β-catenin activator) or GSK3β (Wnt/β-catenin inhibitor). Scale bar; 200 μm

**Figure S4.**
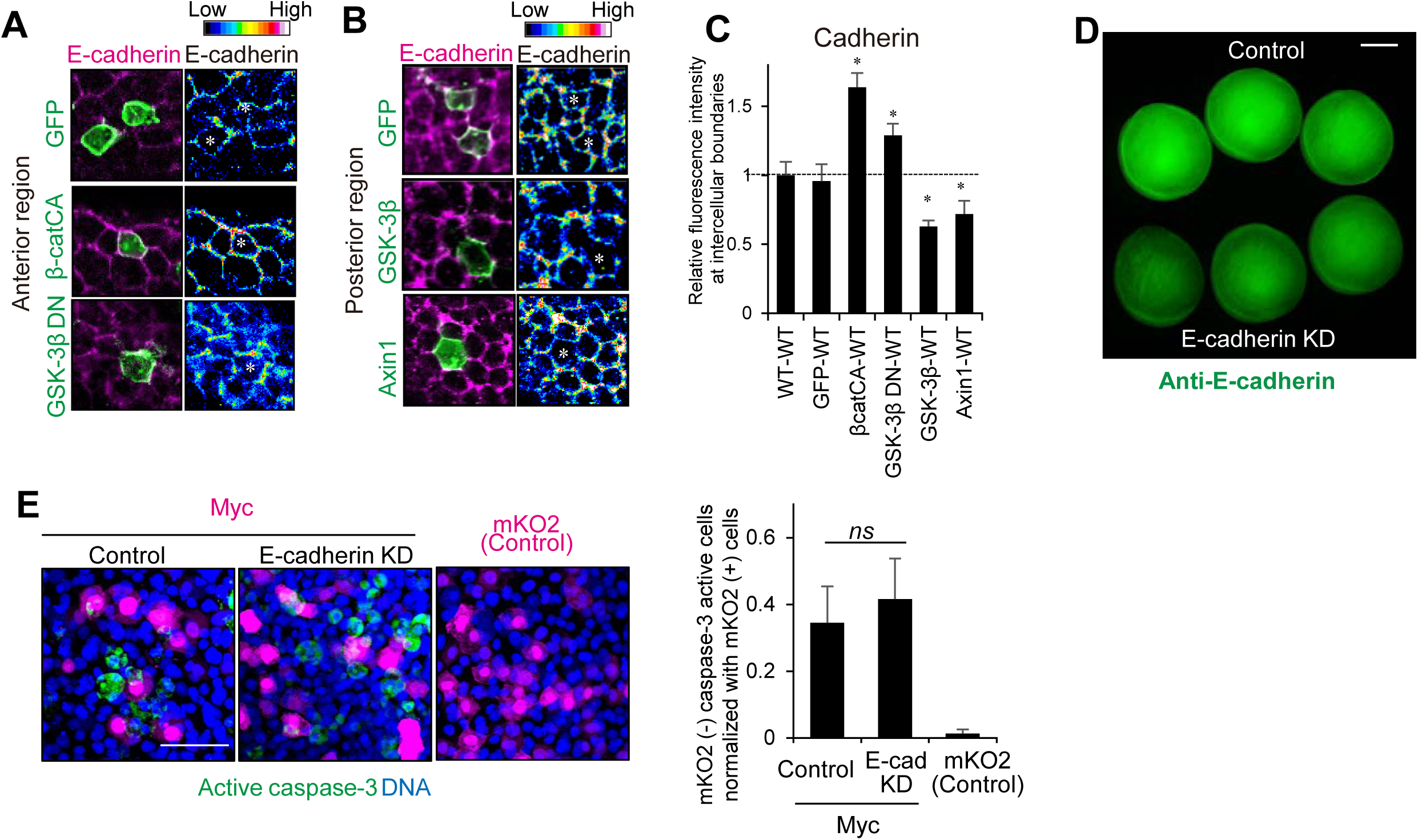
Cadherin mediates Wnt/β-catenin-unfit cell apoptosis, but not Myc-driven cell competition. Related to Figure 4. (A-C) Mosaic introduction of unfit cells with abnormal Wnt/β-catenin activity changes the levels of endogenous E-cadherin. Membrane GFP-expressing cells (GFP), Wnt/β-catenin-activated cells (cells overexpressing βcatCA or GSK3βDN with GFP), or Wnt/β-catenin-inhibited cells (cells overexpressing GSK3β or Axin1 with GFP) were mosaically introduced into zebrafish embryos. Representative confocal fluorescence images in (A) and (B) show GFP^+^ cells and endogenous E-cadherin proteins (magenta and 16 colors of ImageJ) visualized by immunostaining with anti-E-cadherin. The color bar indicates fluorescence intensity from high (red) to low (blue). Wnt/β-catenin-activation increased E-cadherin levels in the anterior region where endogenous E-cadherin levels are low, whereas Wnt/β-catenin-inhibition reduced E-cadherin levels in the posterior region where endogenous E-cadherin levels are high. (C) Fluorescence intensity (means ± SEM) of intercellular E-cadherin staining between GFP^+^ cells and neighboring wild-type cells, normalized to the intercellular fluorescence intensity between wild-type cells in identical each image. Fluorescence intensity in two intercellular areas per cell was measured. \**p* < 0.05. (D) Partial knockdown of E-cadherin (*cdh1*) in zebrafish embryos. Fluorescence image shows anti-E-cadherin immunostaining of embryos injected with 0.3 ng of control MO (upper) or E-cadherin MO (lower). Scale bar, 250 μm. E-cadherin MO reduces the fluorescence signal intensity, suggesting that this MO can reduce endogenous E-cadherin levels and that this E-cadherin antibody recognizes endogenous E-cadherin. (E) High levels of Myc drives cell competition in an E-cadherin-independent manner. Representative confocal fluorescence images show mKO2 (magenta), active caspase-3 (green), and DNA (blue) in 0.3 ng of control MO (control)- or E-cadherin MO-injected (E-cad KD) embryos, which were mosaically introduced with cells overexpressing membrane mKO2 alone (mKO2) or with Myc. Scale bar, 50 μm. Means ± SEM of mKO2-negative caspae-3 active cell numbers normalized with mKO2-positive cells are shown.

**Figure S5.**
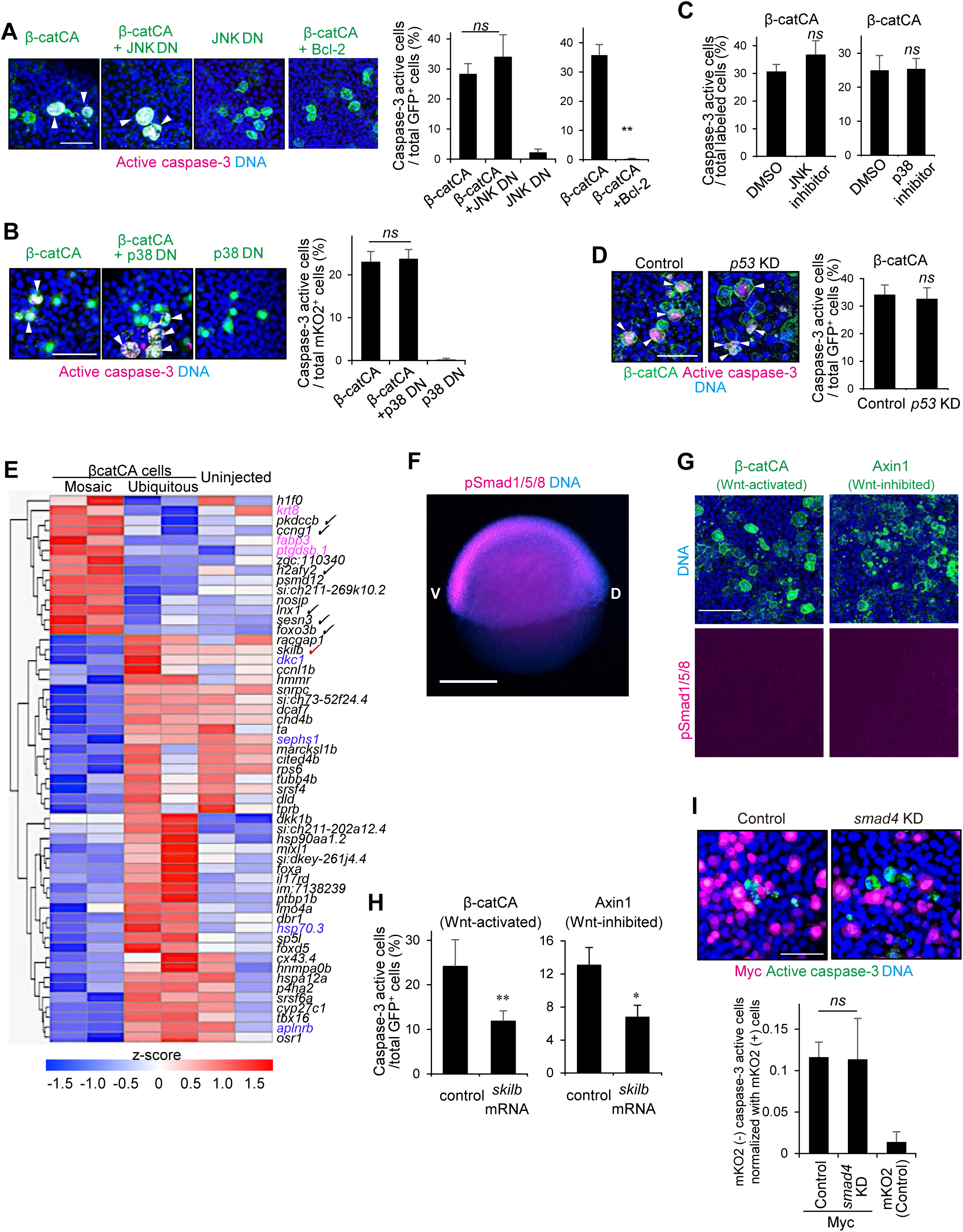
Exploration of the Wnt/β-catenin-unfit cell-killing system. Related to Figure 5. (A) Dominant-negative mutant of JNK (JNK DN) did not block β-catCA-expressing cell apoptosis. Representative confocal fluorescence images show GFP (green), active caspase-3 (magenta), and DNA (blue) in embryos mosaically introduced with cells expressing membrane GFP with β-catCA and JNK DN, as indicated. Means ± SEM of GFP^+^ caspase-3 active cell frequencies are shown. Bcl-2 was used as a positive control. Injection of *bcl2* mRNA (200 pg) efficiently blocked induction of Wnt/β-catenin abnormal-cell apoptosis. (B) Dominant-negative mutant of p38 (p38 DN) did not block apoptosis of β-catCA-expressing cells. Representative confocal fluorescence images show mKO2 (green), active caspase-3 (magenta), and DNA (blue) in embryos mosaically introduced with cells expressing membrane mKO2 with β-catCA, JNK DN, and Bcl-2, as indicated. Means ± SEM of mKO2^+^ caspase-3 active cell frequencies are shown. \*\**p* < 0.01. (C) Treatment with chemical inhibitors against JNK or p38 did not block apoptosis of β-catCA-expressing cells. Embryos were mosaically introduced with cells expressing membrane GFP with β-catCA in the presence of 0.4% DMSO, JNK inhibitor (SP600125 in DMSO, 10 μm), or p38 inhibitor (SB203580 in DMSO, 100 μm). Representative confocal fluorescence images show GFP (green), active caspase-3 (magenta), and DNA (blue) in the embryos. Means ± SEM of GFP^+^ caspase-3 active cell frequencies are shown. Note that treatment with 10 μm SP600125 or 100 μm SB203580 induced morphological defects related to the loss of JNK or p38 (Hsu et al., 2011; Seo et al., 2010), respectively (Y.A. unpublished observations), suggesting that SP600125 and SB203580 blocked endogenous JNK and p38 activities, respectively. (D) Knock-down of p53 did not block apoptosis of β-catCA-expressing cells. Representative fluorescence images show GFP (green), active caspase-3 (magenta), and DNA (blue) of control MO- or *p53* MO-injected embryos mosaically introduced with cells expressing both membrane GFP and β-catCA. Means ± SEM of GFP^+^ (β-catCA) caspase-3 active cell frequencies are shown. (E) Heat map of differentially expressed genes in Mosaic, Ubiquitous, and Uninjected embryos as evaluated by RNA-Seq (n = 2). Check marks: Smad signaling-related genes; Pink and blue genes are positive and negative ROS signaling regulators, respectively. (F, G) BMP-type Smad1/5/8 are not activated in Wnt/β-catenin abnormal cells. As a positive control of immunostaining to phospho-Smad1/5/8, representative fluorescent image show whole-mount immunostaining of anti-phospho-Smad1/5/8 (magenta) and DNA (blue) in zebrafish early embryo at 6 hpf (shield stage) (lateral view with anterior to the top, ventral to the left). V and D indicate ventral and dorsal. Scale bar, 250 μm. (G) Representative confocal fluorescent images show that GFP (green) with β-catCA or Axin1, DNA (blue) and pSmad1/5/8 (magenta). Scale bar, 50 μm. (H) Co-expression of *skilb* blocks apoptosis of β-catCA- or Axin1-expressing cells. Uninjected (control) or *skilb* mRNA (800 pg)-injected embryos were mosaically introduced with cells overexpressing membrane GFP with β-catCA or Axin1 and then caspase-3 activation was detected. Means ± SEM of GFP^+^ caspase-3 active cell frequencies are shown. \*\**p* < 0.01, \**p* < 0.05. (I) Smad4 knock-down does not block Myc-driven cell competition. Representative confocal fluorescence images show mKO2 (magenta), active caspase-3 (green), and DNA (blue) in 10 ng of control MO- or *smad4* MO-injected embryos, which were mosaically introduced with cells overexpressing membrane mKO2 with Myc. Scale bar, 200 μm. Means ± SEM of mKO2-negative caspae-3 active cell numbers normalized with Myc-expressing cells are shown.

**Figure S6.**
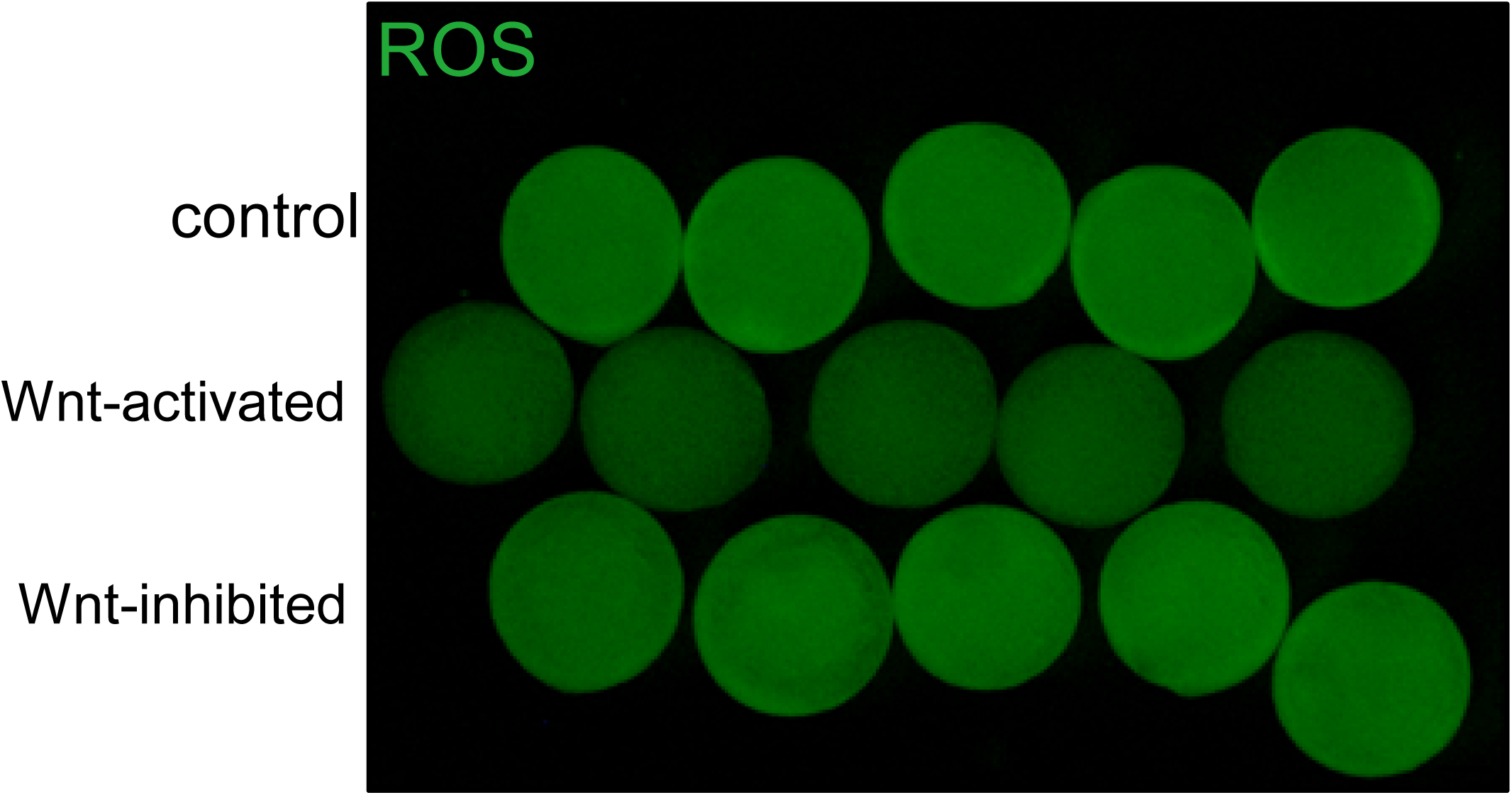
Ubiquitous Wnt/β-catenin activation or inhibition does not activate ROS production. Related to Figure 6. Fluorescence images showing ROS probe (CellRox Green)-stained embryos untreated (top) or treated with 10 μm BIO (Wnt/β-catenin-activator) (middle) or injected with 800 ng of *Dkk1b* mRNA (Wnt/β-catenin-inhibitor) (bottom). Scale bar, 200 μm. Note that ubiquitous Wnt/β-catenin activation slightly reduced endogenous ROS.

**Figure S7.**
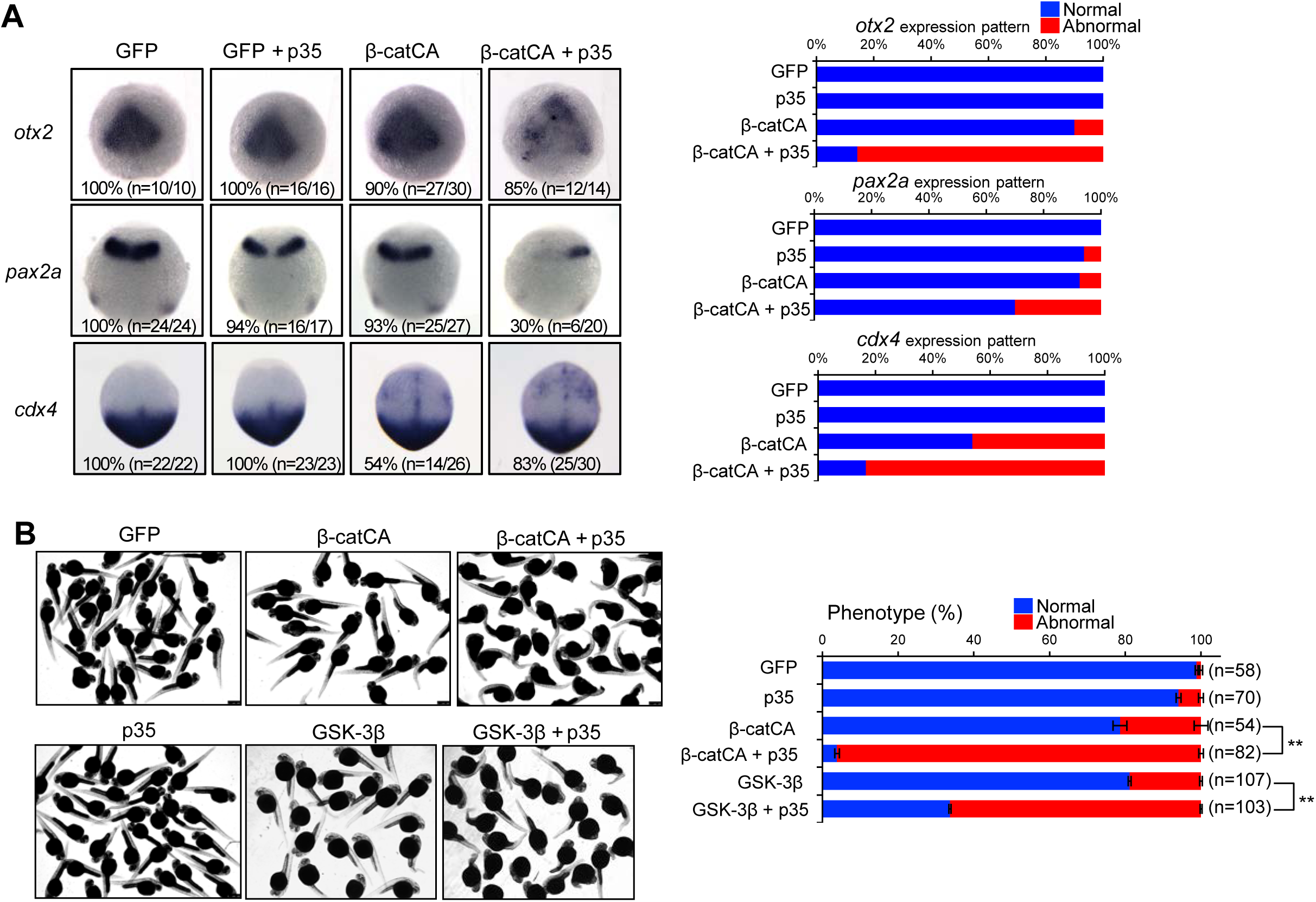
Apoptotic elimination of unfit cells is required for precise AP patterning and morphogenesis. Related to Figure 7. (A) Inhibition of apoptosis in β-catCA-overexpressing cells disturbs AP patterning. Images show whole-mount *in situ* hybridization of *otx2* (marker of presumptive forebrain and midbrain), *pax2a* (marker of presumptive midbrain-hindbrain boundary), and *cdx4* (marker of presumptive spinal cord) in embryos mosaically introduced with cells expressing membrane GFP alone (GFP) or with β-catCA with or without p35. Percentages of embryos displaying abnormal expression patterns are shown. p35 co-expression in artificially introduced β-catCA-expressing cells induced ectopic expression of posterior marker *cdx4* in the anterior tissue and perturbed the expression of anterior markers *otx2* and *pax2a*. (B) Inhibition of apoptosis of β-catCA- or GSK-3β-expressing (Wnt-activated or -inhibited) cells disturbs proper morphogenesis. Embryos artificially introduced with cells overexpressing membrane GFP alone (GFP) or with β-catCA or GSK-3β with or without caspase inhibitor p35. Multi-sample images of 32 hpf larvae. Scale bar, 500 μm. Embryo percentages with normal or abnormal morphogenesis (right). Total analyzed embryo numbers are shown. \*\**p* < 0.01.

**Table S1.**
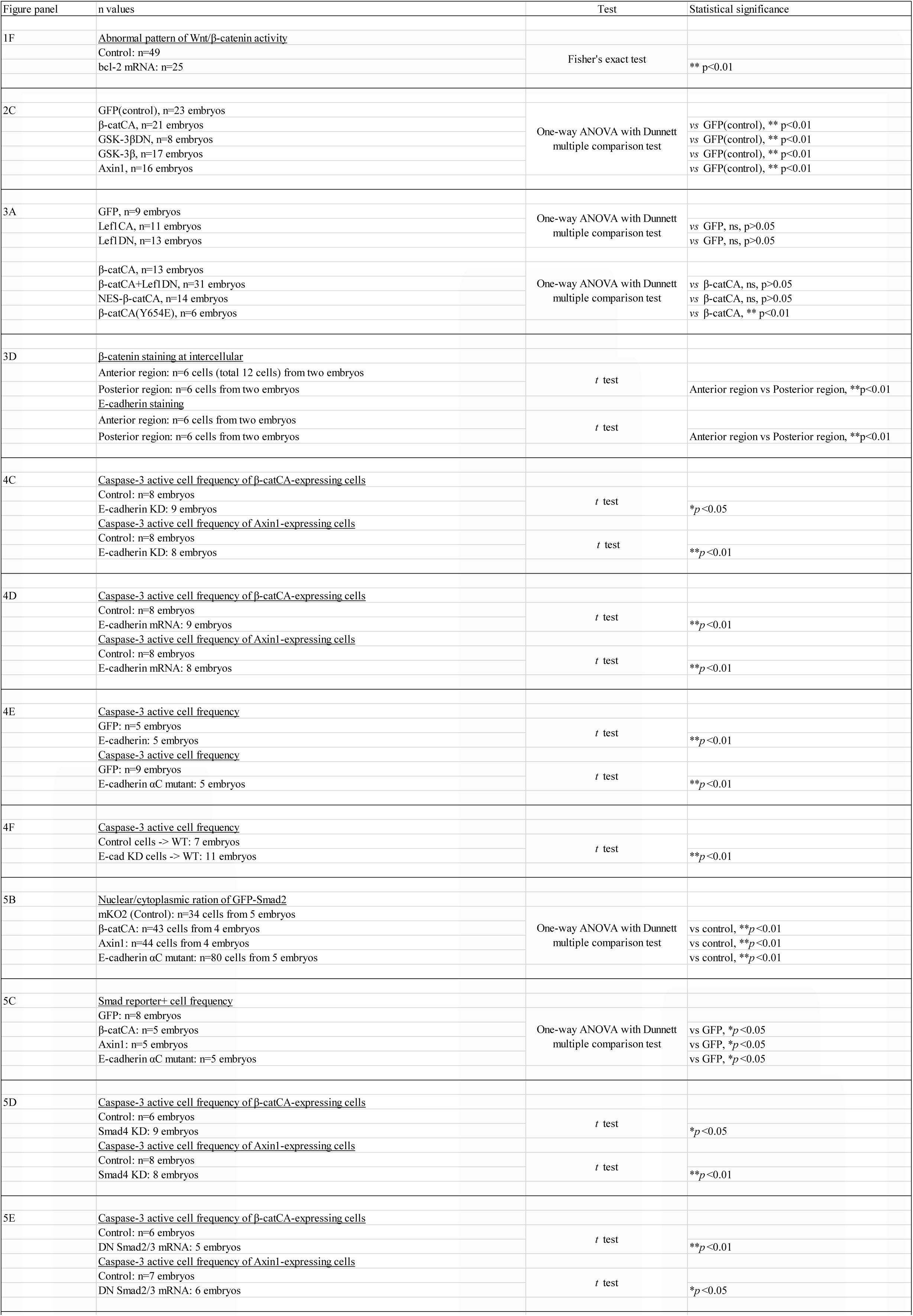
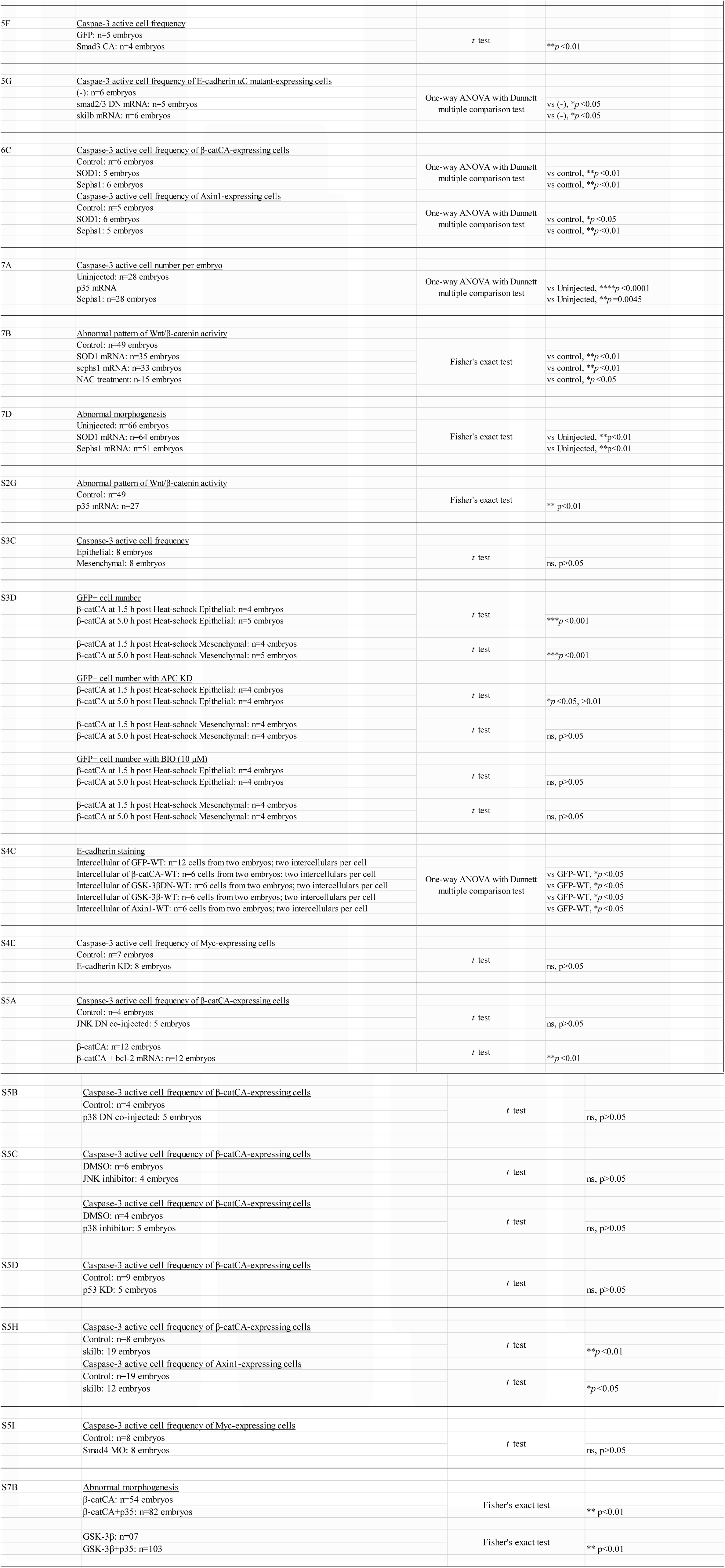
Statistical details and sample number of figures.

**Movie S1. Time-lapse imaging of Wnt/β-catenin signaling activity during zebrafish early embryogenesis**

Wnt/β-catenin activity visualized by ELuc-CP luminescence (merged with bright-field image; left) in Tg(OTM:Eluc-CP) embryo (dorsal view). The luminescence intensity is shown by pseudo-color (16 colors of ImageJ). Bar indicates luminescence intensity from high (red) to low (blue) (center). Images were acquired every 5 min from 8 to 12 hpf. A: anterior, P: posterior. Scale bar, 200 μm. Graph shows time-lapse 3D surface plot of luminescence intensity (right).

**Movie S2. Live imaging of caspase-3 activation in zebrafish embryos**

3D projection of a zebrafish embryo injected with mRNA of the caspase activity fluorescent biosensor/VC3Ai (green) and membrane-tagged mKO2 (magenta) from 7 to 10 hpf. Images were acquired at 1.2-min intervals. The video is displayed at 15 frames/s. Scale bar, 100 μm.

**Movie S3. Behavior of β-catCA-expressing cells in zebrafish early embryos**

3D projection of a mosaic embryos with cells overexpressing membrane GFP with β-catCA (Wnt activator) (green) around 7 hpf (left). The cell membrane was visualized with membrane mKO2 (magenta). Magnified image of boxed area (white line) (right). Images were acquired at 2-min intervals. The video is displayed at 15 frames/s. Scale bar, 100 μm.

**Movie S4. Behavior of GFP-expressing control cells in zebrafish early embryos.**

3D projection of a mosaic embryos with cells overexpressing membrane GFP alone (green) around 7 hpf (left). The cell membrane was visualized with membrane mKO2 (magenta). Magnified image of boxed area (white line) (right). Images were acquired at 2-min intervals. The video is displayed at 15 frames/s. Scale bar, 100 μm.

## Supplemental experimental procedures

### Plasmids

To prepare the luciferase-based reporter plasmid (OTM:ELuc-CP) (Figure S1A), d2EGFP of the OTM:d2EGFP plasmid was replaced with ELuc-PEST (Toyobo, Osaka, Japan) (Shimizu et al., 2012). To enhance destabilization efficiency, the CL1 sequence from pGL4.28 (Promega, Madison, WI) was inserted between ELuc cDNA and the PEST sequence. To prepare heat-shock promoter-driven plasmids, the *hsp70l* promoter was sub-cloned into the pTol2 vector (a gift from Dr. K. Kawakami). Subsequently, membrane-tagged (GAP43-fused) GFP (or mKO2) and T2A were sub-cloned into pTol2-hsp70l promoter plasmids. These plasmids express membrane GFP (or mKO2) alone in response to heat-shock. To generate the plasmids that express membrane GFP (or mKO2) with signaling modulator proteins, PCR-amplified cDNAs encoding signaling modulator proteins were sub-cloned into the downstream site of T2A of pTol2-hsp70l:GFP-T2A plasmids. Wnt/β-catenin signaling activators were as follows: N-terminus truncated mouse β-catenin (β-catCA) (Aberle et al., 1997); NES sequence (PKIα-derived)-conjugated β-catCA (NES-β-catCA); β-catCA Y654E mutant (Huber and Weis, 2001; Roura et al., 1999; van Veelen et al., 2011); dominant-negative mutant of human GSK-3β (GSK-3βDN), in which Arg96 was replaced with Ala (Hagen et al., 2002); constitutively active form of N-terminus-truncated human LRP6 (LRP6CA) (Brennan et al., 2004) (a gift from Dr. A. Kikuchi); and constitutively active human Lef1 mutant (Lef1CA), in which the N-terminus β-catenin-binding region was replaced with the transactivation domain of VP16 (Ishitani et al., 2005). Wnt/β-catenin signaling inhibitors were as follows: human-wild type GSK-3β (a gift from Dr. A. Kikuchi); rat Axin1; dominant-negative form of C-terminus-truncated LRP6 (Tamai et al., 2000) (a gift from Dr. A. Kikuchi); and dominant-negative mutant of human Lef1 (Lef1DN), in which the N-terminus β-catenin binding region was truncated (Yokoyama et al., 2010). Other signaling regulators were as follows: baculovirus-derived p35 (Riedl et al., 2001) (a gift from Dr. G. Salvesen, Addgene plasmid #11808; Cambridge, MA); zebrafish E-cadherin/Cdh1 (a gift from Dr. E. Raz) (Kardash et al., 2010); E-cadherin αC mutant, in which zebrafish E-cadherin N-terminal fragment (1-794 a.a.) lacking the C-terminal β-catenin-binding domain was fused to α-catenin (constructed with reference to mouse E-cadherin αC mutant, described in Nagafuchi et al., 1994); zebrafish *skilb*; dominant-negative zebrafish Smad2 mutant (Smad2 DN), in which Pro446, Ser466, and Ser468 were substituted to His, Ala, and Ala, respectively (Jia et al., 2008); dominant-negative zebrafish Smad3b mutant (Smad3 DN), in which Pro401, Ser421, and Ser423 were substituted to His, Ala, and Ala, respectively (Jia et al., 2008); constitutively active zebrafish Smad3b mutant (Smad3 CA), in which both Ser421 and Ser423 were substituted to Asp; human SOD1 (a gift from Dr. E. Fisher, Addgene #26407) (Stevens et al., 2010); zebrafish Sephs1 (transOMICS tech, Huntsville, AL); human c-Myc (a gift from W. El-Deiry, Addgene #16011) (Ricci et al., 2004); dominant-negative mutant of rat JNK2 (JNK DN), in which the MAPKK-phosphorylation sites <u>Thr</u>-Pro-<u>Tyr</u> were substituted to <u>Ala</u>-Pro-<u>Phe</u> mutants; dominant-negative mutant of *Xenopus* p38β (p38DN), in which an ATP-binding site (Lys53) was substituted to Met; the Rous sarcoma virus *src* (v-Src), which is a constitutively active Src kinase (a gift from Dr. Y. Fujita) (Hunter and Sefon, 1980); and an oncogenic constitutively active mutant of human H-Ras (RasV12), in which Gly12 was substituted to Val (a gift from Dr. A. Yoshimura) (Grand and Owen, 1991). To prepare Krt4P:mCherry, *Gal4* cDNA of the Krt4P:Gal4 plasmid, which expresses Gal4 in epithelia under the control of the keratin4 (Krt4) promoter (Wada et al., 2013), was replaced with mCherry. For mRNA synthesis, cDNAs for signaling proteins were PCR-amplified and cloned into the multi-cloning site of the pCS2p+ vector. Cloned signaling proteins were as follows: human Bcl-2 (a gift from Dr. S. Korsmeyer, Addgene #8768) (Yamamoto et al., 1999), p35, zebrafish E-cadherin-GFP fusion gene (a gift from E. Raz) (Kardash et al., 2010), zebrafish *Dkk1b* (a gift from Dr. M. Hibi) (Hashimoto et al., 2000), GFP-fusion zebrafish Smad2, zebrafish Smad2 DN, zebrafish Smad3a DN, zebrafish Smad3b DN, zebrafish Skilb, human SOD1, and caspase activity fluorescent biosensor/VC3Ai (a gift from Dr. B. Li, Addgene plasmid #78907) (Zhang et al., 2013). Probes for *in situ* hybridization were as follows: *otx2* (Li et al., 1994), *pax2a* (Krauss et al., 1991), *cdx4* (Joly et al., 1992), GFP, Eluc-CP, and firefly luciferase. SBE-luc (pGL4.48[luc2P/SBE/Hygro] Vector) was purchased from Promega.

### Antibodies

Primary antibodies were as follows: anti-E-cadherin (#610181, BD Bioscience, Franklin Lakes, NJ); anti-αTubulin (#T6074, Sigma-Aldrich, St. Louis, MO); mouse anti-GFP (#A-11120, Thermo Fisher, Waltham, MA); rabbit anti-GFP (#A-11122, Thermo Fisher); anti-active caspase-3 (#559565, BD Bioscience); anti-β-catenin (#C7207 Sigma-Aldrich); anti-Sephs1 (ab96542 Abcam); and anti-mKO2 (#M168-3M, MBL, Nagoya, Japan). Secondary antibodies were as follows: AlexaFluor488-conjugated anti-mouse IgG (#A-11029, Invitrogen, Waltham, MA) and anti-rabbit IgG (#A-11034, Invitrogen); AlexaFluor594-conjugated anti-mouse IgG (#A-11032, Invitrogen) and anti-rabbit IgG (#A-11037, Invitrogen); AlexaFluor647-conjugated anti-rabbit IgG (#4414, Cell Signaling Technology, Mountain View, CA).

### TUNEL assay

Embryos were fixed with 4% paraformaldehyde overnight at 4 °C and dechorionated. Embryos were rehydrated using methanol. TUNEL assay was performed with the *In Situ* Cell Death Detection Kit, POD (Roche) according to the manufacturer’s instructions.

### Cell culture, transfection, and reporter gene assay

Neruro-2a neuroblastoma cells were grown in Dulbecco’s modified Eagle’s medium supplemented with 10% fetal bovine serum. Cells were transfected with the expression plasmids using Polyethylenimine MW 25000 (Polysciences, Warrington, PA). Cells were seeded into 35-mm diameter plates and then transfected with OTM:Eluc-CP reporter gene plasmids (500 ng) and the pRL-EF vector (1 ng), which expresses *Renilla* luciferase under the control of the EF-1α promoter, along with the indicated expression vectors. After 48 h, Eluc-CP and *Renilla* luciferase activities were determined using the Dual luciferase assay system (Promega). The pRL-EF vector was used to normalize the transfection efficiency of the luciferase reporters. The mean and standard deviation of duplicates from one of three independent experiments are presented.

### Southern blotting

The tail fin of adult transgenic fish was amputated using surgical scissors and transferred to lysis buffer containing 0.1 μg/μl proteinase K. The sample was incubated overnight at 55 °C, followed by standard ethanol precipitation. Purified genomic DNA samples were digested with restriction endonuclease. Southern blot hybridization was performed using digoxigenin-labeled probe and standard methods.

### mRNA and antisense oligo MO microinjection

Capped mRNA was synthesized using the SP6 mMessage mMachine kit (Ambion, Austin, TX) and purified using Micro Bio-Spin columns (Bio-Rad, Hercules, CA). We injected synthesized mRNA at the one-cell stage of zebrafish embryos.

To perform knock-down experiment in zebrafish embryos, antisense oligo MOs (Gene Tools, Philomath, OR) were injected into one-cell and/or two-cell stage embryos. Standard control morpholino, 5 ′- CCT CTT ACC TCA GTT ACA ATT TAT A-3′; E-cadherin (*cdh1*), 5′-AAA GTC TTA CCT GAA AAA GAA AAA C-3′ (Shimizu et al., 2005); Smad4 (*smad4a*), 5′-AAT CAT ACT CAT CCT TCA CCA TCA T-3′ (Sun et al., 2014); APC, 5′-TAG CAT ACT CTA CCT GTG CTC TTC G-3′ (Nadauld et al., 2004); and p53, 5′-GCG CCA TTG CTT TGC AAG AAT TG-3′ (Robu et al., 2007) were used.

### Transplantation

In the Figure S3E experiment, the embryonic cells from Tg(hsp70l:GFP-T2A-β-catCA) embryos injected with tetramethylrhodamine-conjugated dextran (Molecular Probes, Waltham, MA) were transplanted into 3.3 hpf to 3.7 hpf-stage embryos obtained by crossing heterozygous Tg(hsp70l:GFP-T2A-β-catCA) fish with wild-type fish, and then exposed to heat-shock from 4.3 hpf to 5.3 hpf. The embryos were fixed with 4% paraformaldehyde at 9 hpf, and then immunostained. In the Figure 4F experiment, the embryonic cells from embryos injected with either E-cadherin (*cdh1*) antisense oligo morpholino (1 ng) or standard control morpholino and tetramethylrhodamine-conjugated dextran were transplanted into 3.3 hpf to 3.7 hpf-stage wild-type embryos. The embryos were fixed at 9 hpf and immunostained.

### Chemical inhibitors

JNK inhibitor SP600125 (TOCRIS, Bristol, UK); p38 inhibitor SB203580 (TOCRIS); GSK-3β inhibitor 6-Bromoindirubin-3′-oxime (BIO) (FUJIFILM Wako, Osaka, Japan), and ROS inhibitor N-acetyl-L-cysteine (NAC) (Sigma-Aldrich) were used.

### Western blotting

A total of 30 zebrafish embryos were collected and their chorions and yolks were removed, then embryonic tissues were lysed as previously described (Link et al., 2006). Cell lysates were resolved by sodium dodecyl sulfate-polyacrylamide gel electrophoresis and transferred onto polyvinylidene fluoride membrane (GE Healthcare, Buckinghamshire, UK). Membranes were immunoblotted with the primary antibodies and the bound antibodies were visualized with horseradish-peroxidase-conjugated antibodies against rabbit or mouse IgG (EMD Chemicals Inc., Darmstadt, Germany) using the ChemiLumi-One L Chemiluminescent Kit (Nacalai Tesque, Kyoto, Japan). The band intensities were measured using ImageJ.

### Quantitative reverse transcription (qRT)-PCR

Total RNA from a total 25 zebrafish embryos was purified using TRIzol reagent (Invitrogen) and cDNA was synthesized with the ReverTra Ace qPCR RT Master Mix with gDNA Remover (TOYOBO, Osaka, Japan). qPCR was performed in a Mx3000P QPCR system (Agilent Technologies, Santa Clara, CA) with THUNDERBIRD SYBR qPCR Mix (TOYOBO) and specific primers as follows: *cdh1*; 5′-TCA GTA CAG ACC TCG ACC GGC CAA-3′ (forward), 5′-AAA CAC CAG CAG AGA GTC GTA GG-3′ (Reverse), and *actb1*; 5 ′-TGG ACT TTG AGC AGG AGA TGG GAA-3′ (forward), 5 ′-AAG GTG GTC TCA TGG ATA CCG CAA-3′ (reverse). *actb1* was used as a loading control. qPCR cycling conditions were: 95°C for 1 min, [95°C for 10 s, 60°C for 30 s] (45 cycles), followed by dissociation curve analysis.

### Whole-mount immunostaining

Embryos were fixed with 4% paraformaldehyde in phosphate-buffered saline (PBS) overnight at 4 °C. The dechorionated embryos were washed with 0.5% Triton X-100 (PBST) four times and blocked with 10% fetal bovine serum, 4% Block Ace (Megmilk Snow Brand, Tokyo, Japan), and 1% dimethylsulphoxide (DMSO) in 0.1% PBST for 1 hour. The embryos were incubated with the primary antibodies overnight 4 °C, then washed and incubated with AlexaFluor-conjugated secondary antibodies overnight at 4 °C. Stained embryos were visualized with the M205FA fluorescent stereo-microscope (Leica, Wetzlar, Germany), LSM700 or FV1000 (Olympus) confocal laser-scanning microscope, or Lightsheet Z.1 Light Sheet microscope (Zeiss). Images were prepared and analyzed with ImageJ or Imaris software (Bitplane). For the quantification of membrane β-catenin and E-cadherin levels, the fluorescence intensity in two intercellular areas per cell (n = 6 cells) in each region was measured. Analyzes were performed in duplicate.

### Whole-mount *in situ* hybridization

Whole-mount *in situ* hybridization was performed according to standard protocol. Fluorescence *in situ* hybridization was performed according to previously described protocol (Brend and Holley, 2009). Digoxigenin- or FITC-labeled RNA antisense probes were prepared from plasmids containing *GFP*, *skilb*, *Eluc-CP*, *luciferase*, *otx2*, *pax2a*, and *cdx4*. Images were taken using an M205A stereo-microscope (Leica) and with an FV1000 confocal laser scanning microscope.

### CEL-Seq2 and data analysis

Libraries were constructed following published protocol (Hashimshony et al., 2016) and sequenced using the Illumina HiSeq 1500 (San Diego, CA). The reads were aligned to the zebrafish reference genome (GRCz10) using Bowtie2 software (version 2.3.1) (Langmead and Salzberg, 2012). Read counting was performed per gene with HTSeq (version 0.6.1) (Anders et al., 2015) so that duplicates in unique molecular identifiers were discarded. Using the resulting counts, genes with false discovery rates < 0.1 were extracted as differentially expressed using R library DESeq2 (version 1.10.1) (Love et al., 2014). Smad (Smad4)-target genes were selected using the ChIP-X database (Lachmann et al., 2010) (http://amp.pharm.mssm.edu/lib/chea.jsp).

## REFERENCES

Adachi-Yamada, T., Fujimura-Kamada, K., Nishida, Y., and Matsumoto, K. (1999). Distortion of proximodistal information causes JNK-dependent apoptosis in *Drosophila* wing. Nature 400, 166–169.

Ashe, H.L., and Briscoe, J. (2006). The interpretation of morphogen gradients. Development 133, 385–394.

Brumby, A.M., and Richardson, H.E. (2003). scribble mutants cooperate with oncogenic Ras or Notch to cause neoplastic overgrowth in *Drosophila*. EMBO J. 22, 5769–5779.

Carvalho, L., and Heisenberg, C.-P. (2010). The yolk syncytial layer in early zebrafish development. Trends Cell Biol. 20, 586–592.

Christofori, G., and Semb, H. (1999). The role of the cell-adhesion molecule E-cadherin as a tumour-suppressor gene. Trends Biochem. Sci. 24, 73–76.

Clavería, C., Giovinazzo, G., Sierra, R., and Torres, M. (2013). Myc-driven endogenous cell competition in the early mammalian embryo. Nature 500, 39–44.

de la Cova, C., Senoo-Matsuda, N., Ziosi, M., Wu, D.C., Bellosta, P., Quinzii, C.M., and Johnston, L.A. (2014). Supercompetitor status of *Drosophila* Myc cells requires p53 as a fitness sensor to reprogram metabolism and promote viability. Cell Metab. 19, 470–483.

Derynck, R., Akhurst, R.J., and Balmain, A. (2001). TGF-beta signaling in tumor suppression and cancer progression. Nat. Genet. 29, 117–129.

Di-Gregorio, A., Bowling, S., and Rodriguez, T.A. (2016). Cell competition and its role in the regulation of cell fitness from development to cancer. Dev. Cell 38, 621–634.

Eivers, E., Fuentealba, L.C., and De Robertis, E.M. (2008). Integrating positional information at the level of Smad1/5/8. Curr. Opin. Genet. Dev. 18, 304–310.

Farin, H.F., Jordens, I., Mosa, M.H., Basak, O., Korving, J., Tauriello, D.V.F., de Punder, K., Angers, S., Peters, P.J., Maurice, M.M., et al. (2016). Visualization of a short-range Wnt gradient in the intestinal stem-cell niche. Nature 530, 340–343.

Furumatsu, T., Matsumoto, E., Kanazawa, T., Fujii, M., Lu, Z., Kajiki, R., and Ozaki, T. (2013). Tensile strain increases expression of CCN2 and COL2A1 by activating TGF-β-Smad2/3 pathway in chondrocytic cells. J. Biomech. 46, 1508–1515.

Gaarenstroom, T., and Hill, C.S. (2014). TGF-β signaling to chromatin: how Smad3 regulate transcription during self-renewal and differentiation. Semin. Cell Dev. Biol. 32, 107–118.

Greenlund, L.J., Deckwerth, T.L., and Johnson, E.M. Jr. (1995). Superoxide dismutase delays neuronal apoptosis: a role for reactive oxygen species in programmed neuronal death. Neuron 14, 303–315.

Grzeschik, N.A., Amin, N., Secombe, J., Brumby, A.M., and Richardson, H.E. (2007). Abnormalities in cell proliferation and apico-basal cell polarity are separable in *Drosophila lgl* mutant clones in the developing eye. Dev. Biol. 311, 106–123.

Gu, B.-W., Fan, J.-M., Bessler, M., and Mason, P.J. (2011). Accelerated hematopoietic stem cell aging in a mouse model of dyskeratosis congenita responds to antioxidant treatment. Aging Cell 10, 338–348.

Hagenbuchner, J., Kuznetsov, A., Hermann, M., Hausott, B., Obexer, P., and Ausserlechner, M.J. (2012). FOXO3-induced reactive oxygen species are regulated by BCL2L11 (Bim) and SESN3. J. Cell. Sci. 125, 1191–1203.

Halbleib, J.M., and Nelson, W.J. (2006). Cadherins in development: cell adhesion, sorting, and tissue morphogenesis. Genes Dev. 20, 3199–3214.

Huang, D., Liu, Y., Huang, Y., Xie, Y., Shen, K., Zhang, D., and Mou, Y. (2014). Mechanical compression upregulates MMP9 through SMAD3 but not SMAD2 modulation in hypertrophic scar fibroblasts. Connect. Tissue Res. 55, 391–396.

Huber, A.H., and Weis, W.I. (2001). The structure of the beta-catenin/E-cadherin complex and the molecular basis of diverse ligand recognition by beta-catenin. Cell 105, 391–402.

Igaki, T. (2009). Correcting developmental errors by apoptosis: lessons from *Drosophila* JNK signaling. Apoptosis 14, 1021–1028.

Johnston, L.A., Prober, D.A., Edgar, B.A., Eisenman, R.N., and Gallant, P. (1999). *Drosophila* myc regulates cellular growth during development. Cell 98, 779–790.

Kinzler, K.W., Nilbert, M.C., Su, L.K., Vogelstein, B., Bryan, T.M., Levy, D.B., Smith, K.J., Preisinger, A.C., Hedge, P., and McKechnie, D. (1991). Identification of *FAP* locus genes from chromosome 5q21. Science 253, 661–665.

Kunnen, S.J., Leonhard, W.N., Semeins, C., Hawinkels, L.J.A.C., Poelma, C., Ten Dijke, P., Bakker, A., Hierck, B.P., and Peters, D.J.M. (2017). Fluid shear stress-induced TGF-β/ALK5 signaling in renal epithelial cells is modulated by MEK1/2. Cell. Mol. Life Sci. 74, 2283–2298.

Lachmann, A., Xu, H., Krishnan, J., Berger, S.I., Mazloom, A.R., and Ma’ayan, A. (2010). ChEA: transcription factor regulation inferred from integrating genome-wide ChIP-X experiments. Bioinformatics 26, 2438–2444.

Link, V., Shevchenko, A., and Heisenberg, C.-P. (2006). Proteomics of early zebrafish embryos. BMC Dev. Biol. 6, 1.

Liu, R.-M., and Desai, L.P. (2015). Reciprocal regulation of TGF-β and reactive oxygen species: A perverse cycle for fibrosis. Redox Biol. 6, 565–577.

Mathew, J., Galarneau, L., Loranger, A., Gilbert, S., and Marceau, N. (2008). Keratin-protein kinase C interaction in reactive oxygen species-induced hepatic cell death through mitochondrial signaling. Free Radic. Biol. Med. 45, 413–424.

Merino, M.M., Levayer, R., and Moreno, E. (2016). Survival of the fittest: essential roles of cell competition in development, aging, and cancer. Trends Cell Biol. 26, 776–788.

Morata, G., and Ripoll, P. (1975). Minutes: mutants of drosophila autonomously affecting cell division rate. Dev. Biol. 42, 211–221.

Moreno, E., Basler, K., and Morata, G. (2002). Cells compete for decapentaplegic survival factor to prevent apoptosis in *Drosophila* wing development. Nature 416, 755–759.

Nagafuchi, A., and Takeichi, M. (1989). Transmembrane control of cadherin-mediated cell adhesion: a 94 kDa protein functionally associated with a specific region of the cytoplasmic domain of E-cadherin. Cell Regul. 1, 37–44.

Nishisho, I., Nakamura, Y., Miyoshi, Y., Miki, Y., Ando, H., Horii, A., Koyama, K., Utsunomiya, J., Baba, S., and Hedge, P. (1991). Mutations of chromosome 5q21 genes in FAP and colorectal cancer patients. Science 253, 665–669.

Norman, M., Wisniewska, K.A., Lawrenson, K., Garcia-Miranda, P., Tada, M., Kajita, M., Mano, H., Ishikawa, S., Ikegawa, M., Shimada, T., et al. (2012). Loss of Scribble causes cell competition in mammalian cells. J. Cell. Sci. 125, 59–66.

Petersen, C.P., and Reddien, P.W. (2009). Wnt signaling and the polarity of the primary body axis. Cell 139, 1056–1068.

Roura, S., Miravet, S., Piedra, J., Garcia de Herreros, A., and Dunach, M. (1999). Regulation of E-cadherin/Catenin association by tyrosine phosphorylation. J. Biol. Chem. 274, 36734–36740.

Saha, S., Ji, L., de Pablo, J.J., and Palecek, S.P. (2008). TGFbeta/Activin/Nodal pathway in inhibition of human embryonic stem cell differentiation by mechanical strain. Biophys. J. 94, 4123–4133.

Sancho, M., Di-Gregorio, A., George, N., Pozzi, S., Sánchez, J.M., Pernaute, B., and Rodríguez, T.A. (2013). Competitive interactions eliminate unfit embryonic stem cells at the onset of differentiation. Dev. Cell 26, 19–30.

Schier, A.F., and Talbot, W.S. (2005). Molecular genetics of axis formation in zebrafish. Annu. Rev. Genet. 39, 561–613.

Shen, Y.-H., Song, G.-X., Liu, Y.-Q., Sun, W., Zhou, L.-J., Liu, H.-L., Yang, R., Sheng, Y.-H., Qian, L.-M., and Kong, X.-Q. (2012). Silencing of FABP3 promotes apoptosis and induces mitochondrion impairment in embryonic carcinoma cells. J. Bioenerg. Biomembr. 44, 317–323.

Shi, K., Zhang, X., Lei, Y., Tan, J., Xu, Z., and Zheng, Y. (2016). HSP70 alleviates PC-12 cell oxidative damage caused by ROS through activation of Wnt/beta-catenin signaling pathway in spinal cord injury. Int. J. Clin. Exp. Pathol. 9, 12581–12587.

Shimizu, N., Kawakami, K., and Ishitani, T. (2012). Visualization and exploration of Tcf/Lef function using a highly responsive Wnt/β-catenin signaling-reporter transgenic zebrafish. Dev. Biol. 370, 71–85.

Steinberg, M.S., and Takeichi, M. (1994). Experimental specification of cell sorting, tissue spreading, and specific spatial patterning by quantitative differences in cadherin expression. Proc. Natl. Acad. Sci. U. S. A. 91, 206–209.

Stoick-Cooper, C.L., Weidinger, G., Riehle, K.J., Hubbert, C., Major, M.B., Fausto, N., and Moon, R.T. (2007). Distinct Wnt signaling pathways have opposing roles in appendage regeneration. Development 134, 479–489.

Sugita, K., Ikenouchi-Sugita, A., Nakayama, Y., Yoshioka, H., Nomura, T., Sakabe, J.-I., Nakahigashi, K., Kuroda, E., Uematsu, S., Nakamura, J., et al. (2013). Prostaglandin E_2_ is critical for the development of niacin-deficiency-induced photosensitivity via ROS production. Sci. Rep. 3, 2973.

Takeichi, M. (1991). Cadherin cell adhesion receptors as a morphogenetic regulator. Science 251, 1451–1455.

Than, A., Zhang, X., Leow, M.K.-S., Poh, C.L., Chong, S.K., and Chen, P. (2014). Apelin attenuates oxidative stress in human adipocytes. J. Biol. Chem. 289, 3763–3774.

Tobe, R., Carlson, B.A., Huh, J.H., Castro, N.P., Xu, X.-M., Tsuji, P.A., Lee, S.-G., Bang, J., Na, J.-W., Kong, Y.-Y., et al. (2016). Selenophosphate synthetase 1 is an essential protein with roles in regulation of redox homoeostasis in mammals. Biochem. J. 473, 2141–2154.

van Veelen, W., Le, N.H., Helvensteijn, W., Blonden, L., Theeuwes, M., Bakker, E.R.M., Franken, P.F., van Gurp, L., Meijlink, F., van der Valk, M.A., et al. (2011). β-catenin tyrosine 654 phosphorylation increases Wnt signalling and intestinal tumorigenesis. Gut 60, 1204–1212.

Vincent, J.-P., Kolahgar, G., Gagliardi, M., and Piddini, E. (2011). Steep differences in wingless signaling trigger Myc-independent competitive cell interactions. Dev. Cell 21, 366–374.

Wolpert, L. (1969). Positional information and the spatial pattern of cellular differentiation. J. Theor. Biol. 25, 1–47.

Wong, M.H., Rubinfeld, B., and Gordon, J.I. (1998). Effects of forced expression of an NH_2_-terminal truncated beta-Catenin on mouse intestinal epithelial homeostasis. J. Cell Biol. 141, 765–777.

Xiong, F., Tentner, A.R., Huang, P., Gelas, A., Mosaliganti, K.R., Souhait, L., Rannou, N., Swinburne, I.A., Obholzer, N.D., Cowgill, P.D., et al. (2013). Specified neural progenitors sort to form sharp domains after noisy Shh signaling. Cell 153, 550–561.

Yamamoto, M., Ohsawa, S., Kunimasa, K., and Igaki, T. (2017). The ligand Sas and its receptor PTP10D drive tumour-suppressive cell competition. Nature 542, 246–250.

## Supplemental references

Aberle, H., Bauer, A., Stappert, J., Kispert, A., and Kemler, R. (1997). Beta-catenin is a target for the ubiquitin-proteasome pathway. EMBO J. 16, 3797–3804.

Anders, S., Pyl, P.T., and Huber, W. (2015). HTSeq--a Python framework to work with high-throughput sequencing data. Bioinformatics 31, 166–169.

Asakawa, K., Suster, M.L., Mizusawa, K., Nagayoshi, S., Kotani, T., Urasaki, A., Kishimoto, Y., Hibi, M., and Kawakami, K. (2008). Genetic dissection of neural circuits by Tol2 transposon-mediated *Gal4* gene and enhancer trapping in zebrafish. Proc. Natl. Acad. Sci. U. S. A. 105, 1255–1260.

Brend, T., and Holley, S.A. (2009). Zebrafish whole mount high-resolution double fluorescent in situ hybridization. J. Vis. Exp. pii=1229.

Brennan, K., Gonzalez-Sancho, J.M., Castelo-Soccio, L.A., Howe, L.R., and Brown, A.M.C. (2004). Truncated mutants of the putative Wnt receptor LRP6/Arrow can stabilize beta-catenin independently of Frizzled proteins. Oncogene 23, 4873–4884.

Grand, R.J., and Owen, D. (1991). The biochemistry of ras p21. Biochem. J. 279, 609–631.

Hagen, T., Di Daniel, E., Culbert, A.A., and Reith, A.D. (2002). Expression and characterization of GSK-3 mutants and their effect on beta-catenin phosphorylation in intact cells. J. Biol. Chem. 277, 23330–23335.

Hashimoto, H., Itoh, M., Yamanaka, Y., Yamashita, S., Shimizu, T., Solnica-Krezel, L., Hibi, M., and Hirano, T. (2000). Zebrafish Dkk1 functions in forebrain specification and axial mesendoderm formation. Dev. Biol. 217, 138–152.

Hashimshony, T., Senderovich, N., Avital, G., Klochendler, A., de Leeuw, Y., Anavy, L., Gennert, D., Li, S., Livak, K.J., Rozenblatt-Rosen, O., et al. (2016). CEL-Seq2: sensitive highly-multiplexed single-cell RNA-Seq. Genome Biol. 17, 77.

Hsu, R.-J., Lin, C.-C., Su, Y.-F., and Tsai, H.-J. (2011). Dickkopf-3-related gene regulates the expression of zebrafish *myf5* gene through phosphorylated p38a-dependent Smad4 activity. J. Biol. Chem. 286, 6855–6864.

Huber, A.H., and Weis, W.I. (2001). The structure of the betβ-catenin/E-cadherin complex and the molecular basis of diverse ligand recognition by betβ-catenin. Cell 105, 391–402.

Hunter, T., and Sefton, B.M. (1980). Transforming gene product of Rous sarcoma virus phosphorylates tyrosine. Proc. Natl. Acad. Sci. U. S. A. 77, 1311–1315.

Ishitani, T., Matsumoto, K., Chitnis, A.B., and Itoh, M. (2005). Nrarp functions to modulate neural-crest-cell differentiation by regulating LEF1 protein stability. Nat. Cell Biol. 7, 1106–1112.

Jia, S., Ren, Z., Li, X., Zheng, Y., and Meng, A. (2008). Smad2 and smad3 are required for mesendoderm induction by transforming growth factor-beta/nodal signals in zebrafish. J. Biol. Chem. 283, 2418–2426.

Joly, J.S., Maury, M., Joly, C., Duprey, P., Boulekbache, H., and Condamine, H. (1992). Expression of a zebrafish caudal homeobox gene correlates with the establishment of posterior cell lineages at gastrulation. Differentiation 50, 75–87.

Kardash, E., Reichman-Fried, M., Maître, J.-L., Boldajipour, B., Papusheva, E., Messerschmidt, E.-M., Heisenberg, C.-P., and Raz, E. (2010). A role for Rho GTPases and cell-cell adhesion in single-cell motility in vivo. Nat. Cell Biol. 12, 47–53, supp1-11.

Krauss, S., Johansen, T., Korzh, V., and Fjose, A. (1991). Expression pattern of zebrafish *pax* genes suggests a role in early brain regionalization. Nature 353, 267–270.

Lachmann, A., Xu, H., Krishnan, J., Berger, S.I., Mazloom, A.R., and Ma–ayan, A. (2010). ChEA: transcription factor regulation inferred from integrating genome-wide ChIP-X experiments. Bioinformatics 26, 2438–2444.

Langmead, B., and Salzberg, S.L. (2012). Fast gapped-read alignment with Bowtie 2. Nat. Methods 9, 357–359.

Li, Y., Allende, M.L., Finkelstein, R., and Weinberg, E.S. (1994). Expression of two zebrafish orthodenticle-related genes in the embryonic brain. Mech. Dev. 48, 229–244.

Love, M.I., Huber, W., and Anders, S. (2014). Moderated estimation of fold change and dispersion for RNA-seq data with DESeq2. Genome Biol. 15, 550.

Nadauld, L.D., Sandoval, I.T., Chidester, S., Yost, H.J., and Jones, D.A. (2004). Adenomatous polyposis coli control of retinoic acid biosynthesis is critical for zebrafish intestinal development and differentiation. J. Biol. Chem. 279, 51581–51589.

Nagafuchi, A., Ishihara, S., and Tsukita, S. (1994). The roles of catenins in the cadherin-mediated cell adhesion: functional analysis of E-cadherin-alpha catenin fusion molecules. J. Cell Biol. 127, 235–245.

Ricci, M.S., Jin, Z., Dews, M., Yu, D., Thomas-Tikhonenko, A., Dicker, D.T., and El-Deiry, W.S. (2004). Direct repression of FLIP expression by c-myc is a major determinant of TRAIL sensitivity. Mol. Cell. Biol. 24, 8541–8555.

Riedl, S.J., Renatus, M., Snipas, S.J., and Salvesen, G.S. (2001). Mechanism-based inactivation of caspases by the apoptotic suppressor p35. Biochemistry 40, 13274–13280.

Robu, M.E., Larson, J.D., Nasevicius, A., Beiraghi, S., Brenner, C., Farber, S.A., and Ekker, S.C. (2007). p53 activation by knockdown technologies. PLoS Genet. 3, e78.

Seo, J., Asaoka, Y., Nagai, Y., Hirayama, J., Yamasaki, T., Namae, M., Ohata, S., Shimizu, N., Negishi, T., Kitagawa, D., et al. (2010). Negative regulation of wnt11 expression by Jnk signaling during zebrafish gastrulation. J. Cell. Biochem. 110, 1022–1037.

Shimizu, T., Yabe, T., Muraoka, O., Yonemura, S., Aramaki, S., Hatta, K., Bae, Y.-K., Nojima, H., and Hibi, M. (2005). E-cadherin is required for gastrulation cell movements in zebrafish. Mech. Dev. 122, 747–763.

Stevens, J.C., Chia, R., Hendriks, W.T., Bros-Facer, V., van Minnen, J., Martin, J.E., Jackson, G.S., Greensmith, L., Schiavo, G., and Fisher, E.M.C. (2010). Modification of superoxide dismutase 1 (SOD1) properties by a GFP tag--implications for research into amyotrophic lateral sclerosis (ALS). PLoS One 5, e9541.

Sun, Y., Tseng, W.-C., Fan, X., Ball, R., and Dougan, S.T. (2014). Extraembryonic signals under the control of MGA, Max, and Smad4 are required for dorsoventral patterning. Dev. Cell 28, 322–334.

Tamai, K., Semenov, M., Kato, Y., Spokony, R., Liu, C., Katsuyama, Y., Hess, F., Saint-Jeannet, J.P., and He, X. (2000). LDL-receptor-related proteins in Wnt signal transduction. Nature 407, 530–535.

Wada, H., Ghysen, A., Asakawa, K., Abe, G., Ishitani, T., and Kawakami, K. (2013). Wnt/Dkk negative feedback regulates sensory organ size in zebrafish. Curr Biol 23, 1559–1565.

Yamamoto, K., Ichijo, H., and Korsmeyer, S.J. (1999). BCL-2 is phosphorylated and inactivated by an ASK1/Jun N-terminal protein kinase pathway normally activated at G(2)/M. Mol. Cell. Biol. 19, 8469–8478.

Yokoyama, N.N., Pate, K.T., Sprowl, S., and Waterman, M.L. (2010). A role for YY1 in repression of dominant negative LEF-1 expression in colon cancer. Nucleic Acids Res. 38, 6375–6388.

Zhang, J., Wang, X., Cui, W., Wang, W., Zhang, H., Liu, L., Zhang, Z., Li, Z., Ying, G., Zhang, N., et al. (2013). Visualization of caspase-3-like activity in cells using a genetically encoded fluorescent biosensor activated by protein cleavage. Nat. Commun. 4, 2157.

